# Identifying climatic drivers of hybridization in Heuchereae (Saxifragaceae)

**DOI:** 10.1101/2022.08.24.505154

**Authors:** R.A. Folk, M.L. Gaynor, N.J. Engle-Wrye, B.C. O’Meara, P.S. Soltis, D.E. Soltis, R.P. Guralnick, S.A. Smith, C.J. Grady, Y. Okuyama

**Author notes:** co-first authors (R.A. Folk, M.L. Gaynor).

## Abstract

Applications of molecular phylogenetic approaches have uncovered evidence of hybridization across numerous clades of life, yet the environmental factors responsible for driving opportunities for hybridization remain obscure. Verbal models implicating geographic range shifts that brought species together during the Pleistocene have often been invoked, but quantitative tests using paleoclimatic data are needed to validate these models. Here, we produce a phylogeny for Heuchereae, a clade of 15 genera and 83 species in Saxifragaceae, with complete sampling of recognized species, using 277 nuclear loci and nearly complete chloroplast genomes. We then employ an improved framework with a coalescent simulation approach to test and ultimately confirm previous hybridization hypotheses and identify one new intergeneric hybridization event. Focusing on the North American distribution of Heuchereae, we introduce and implement a newly developed approach to reconstruct potential past distributions for ancestral lineages across all species in the clade and across a paleoclimatic record extending from the late Pliocene. Time calibration based on both nuclear and chloroplast trees recovers a mid- to late-Pleistocene date for most inferred hybridization events, a timeframe concomitant with repeated geographic range restriction into overlapping refugia. Our results indicate an important role for past episodes of climate change, and the contrasting responses of species with differing ecological strategies, in generating novel patterns of range contact among plant communities and therefore new opportunities for hybridization.

## Introduction

Hybridization has long seized the attention of systematists, but a recent appreciation of the magnitude of its importance in the tree of life (reviewed in, among others, Arnold 1997; Soltis and Soltis 2009; Mallet et al. 2016; Payseur and Rieseberg 2016; Folk et al. 2018b) has been inspired by recent advances in genome-scale data collection (recent reviews: McCormack et al. 2013; Jones and Good 2016; McKain et al. 2018; Andermann et al. 2019) and statistical methods (an incomplete but representative list: Huson and Bryant 2006; Joly et al. 2009b; Meng and Kubatko 2009; Yu et al. 2011; Pease and Hahn 2015; Kubatko and Chifman 2019). One of the most significant contributions of molecular data, from allozymes to genomes, has been a recognition of the sheer frequency of hybridization across the tree of life (Folk et al. 2018a), many instances of which were surprises in groups not traditionally thought to undergo hybridization (Garrigan et al. 2012; Liu et al. 2015). Despite substantial progress, much of this work has been devoted to the head and tail of the problem: that is, either to the detection of hybrids (the first step to studying them) or to assessing their adaptive significance (among their furthest downstream implications). We know enough about hybrids now to realize that they truly do lurk “under every bush” (Anderson 1949, p. 101) and that their impact can be significant (Arnold 1997; Soltis and Soltis 2009; Payseur and Rieseberg 2016; Folk et al. 2018b). While our knowledge of the frequency, distribution, and implications of hybridization is rapidly increasing, the external forces that drive hybridization and that may be partly responsible for the remarkable variation in the frequency and consequences of hybridization across taxa and space are still uncertain (reviewed in Folk et al. 2018b). Hybridization’s “controlling factors” (Wiegand 1935) remain obscure.

Despite a limited empirical understanding of the drivers of hybridization, there is a rich body of theory about the interaction of hybridization and the environment, much of it from the botanical literature of the mid-20th century. Most influential has been the hypothesis of Anderson (Anderson 1948; 1949: chapter 2) that hybridization, and specifically the permanent establishment of hybrid progeny, is associated with “hybridized” habitat, that is, habitat intermediate between the parents. Anderson argued that such intermediate habitats are crucial to the initial survival of hybrids with recombined niche traits, and that the generation of such hybridized habitats is primarily due to ecological disturbance. Under the heading of ecological disturbance, Anderson explicitly analogized changes to habitats of anthropogenic origin (Wiegand 1935) and those due to historical climatic origin (Anderson and Stebbins 1954).

Anderson’s disturbance hypothesis was influential (Arnold 2016), and its invocation has been frequent in the literature, but studies have been largely restricted to verbal models (Dobeš et al. 2004; Edwards et al. 2006; Melo-Ferreira et al. 2007; Dixon et al. 2009; Joly et al. 2009a; Majure et al. 2012; Guest and Allen 2014; Sun et al. 2014; López-Alvarez et al. 2015; Klein and Kadereit 2016; Marques et al. 2016; Villanea and Schraiber 2019). In many cases, these scenarios were inspired by strong inferences from molecular data of evidence for hybridization between putative parents that are today widely allopatric (reviewed in Folk et al. 2018b). While valuable, these hypotheses have generally relied primarily on time calibration dating hybridization to the Pleistocene epoch, but without explicit quantitative tests of the impact that past climate change may have had on species distributions (Folk et al. 2018b; also reviewed in Arnold 2016).

Ancestral niche reconstruction (alternatively, “phyloclimatic” modeling; Yesson and Culham 2006) provides an excellent framework for identifying past environmental drivers of hybridization. This set of approaches seeks to combine environmental information from present-day species, often in the form of ecological niche models, with ancestral reconstruction methods to reconstruct past distributions. These reconstructions enable analysis of past climate suitability using paleoclimate data, which are increasingly available with high spatial resolution (Fick and Hijmans 2017; Brown et al. 2018). In its application to hybridization, ancestral niche reconstruction can generate predictions about past ecological niche and the past distribution of suitable habitat, and therefore can directly test the disturbance hypothesis and its prediction of greater overlap and opportunities for interspecific gene flow during past climate change. Perhaps the most conspicuous and most commonly invoked example of a climatic enabler of hybridization would be the Pleistocene period, “the most revolutionary event in the history of the northern continents” (Anderson and Stebbins 1954).

Testing the disturbance hypothesis requires a study system demonstrating hybridization events that span the time period of interest with appropriate evolutionary replication (Folk et al. 2018b). Heuchereae, one of the ten recognized tribes of the plant family Saxifragaceae and a recent radiation of late Miocene origin with 83 currently known species in 15 recognized genera (Folk et al. 2021), meets these requirements. As one of the first groups for which chloroplast capture was reported, Heuchereae remains among the most prolific examples of hybridization in flowering plants, with upwards of a dozen clear examples of interspecific and intergeneric gene flow occurring among and between at least seven of 15 genera: *Conimitella, Heuchera*, *Lithophragma, Mitella, Ozomelis*, *Pectiantia, Tiarella, Tellima* (Fernald 1906; Soltis et al. 1991; Soltis and Kuzoff 1995; Kuzoff et al. 1999; Okuyama et al. 2012; Folk et al. 2017; Liu et al. 2020). The reproductive biology of the group is characterized by weak post-zygotic isolation mechanisms, and artificial hybridization has been demonstrated to be possible among and within many genera (Rosendahl et al. 1936; Leedy 1943; Taylor 1965; Spongberg 1972; Wells 1979; Rabe and Soltis 1999; Okuyama and Kato 2009). Similarly, the broad familiarity of this clade in landscaping is, in part, due to vigorous artificial hybrids among often distantly related species (Heims et al. 2005; Oliver et al. 2006). While polyploidy occurs in Heuchereae, it is sporadic (Folk and Freudenstein 2014) with autopolyploidy thought to be the dominant process in North American species (Soltis and Rieseberg 1986; Ness et al. 1989; Wolf et al. 1989; Folk et al. 2017). The separate occurrence of polyploidization and (homoploid) hybridization therefore yields the opportunity to study the latter process in isolation from the potentially confounding effect of the former (Folk et al. 2018a). In nature, pre-zygotic isolation through habitat and pollinator specialization is thought to form the primary barrier to interspecific gene flow (Rosendahl et al. 1936; Wells 1984; Okamoto et al. 2015; Friberg et al. 2016, 2019; Folk et al. 2021).

As with many other plant groups (Ellstrand et al. 1996; Folk et al. 2018a; Mitchell and Whitney 2021), the reasons for frequent hybridization in Heuchereae remain obscure. While few intrinsic barriers to interspecific gene flow exist in Heuchereae, that is true of many taxa that rarely hybridize in nature, suggesting that external factors, currently poorly understood, may be responsible for the profound disparities in the frequency of hybridization across organismal groups (Ellstrand et al. 1996). One potential explanation, at least for some lineages, is Anderson’s disturbance hypothesis, which would place at the forefront differing ecological histories among clades of life. Previous work (Folk et al. 2018b) identified Pleistocene cooling as a driver of hybridization opportunities in a single pair of focal taxa of Heuchereae, *Heuchera* subsect. *Elegantes* and *Mitella*. Here, we implement new approaches to expand upon this study, testing whether historical climatic changes that promote range overlap are predictive of hybridization. To achieve this historical reconstruction, we first develop a phylogeny of all described species to identify possible cases of hybridization, confirming chloroplast capture events evident from the comparison of nuclear and chloroplast genome histories using greatly improved taxonomic sampling. We then apply a novel method of ancestral niche reconstruction that can capture the shape of niche tolerances without imposing particular assumptions on its distribution, using this to develop a high-resolution history of ancestral temperature niche and potential suitable habitat in North America over much of the evolution of Heuchereae from the mid-Pliocene to the present.

## Methods

### DNA extraction and sequencing

Samples were chosen to completely represent all species-level diversity in Heuchereae. This effort includes the genera *Asimitellaria, Bensoniella, Brewerimitella, Conimitella, Elmera, Heuchera, Lithophragma, Mitella, Mitellastra, Ozomelis, Pectiantia, Spuriomitella, Tellima, Tiarella,* and *Tolmiea*, totaling 83 species currently recognized, as well as 25/29 subspecific taxa recognized in the most recent genus-wide treatment of the large genus *Heuchera* (Folk 2015). Our coverage of the clade represents a substantial improvement over previous studies, which were hampered in part by the difficulty of obtaining several microendemics (Soltis et al. 1991; Soltis and Kuzoff 1995; Okuyama et al. 2012; Folk and Freudenstein 2014; Folk et al. 2017, 2018b, 2021). For this study, sequence data for 54 new accessions (summarized in Supplementary Table S1) were generated following Folk et al. (2015) to increase species and subspecific taxon representation. Briefly, whole genomic DNAs were isolated from fresh, silica-dried, or herbarium leaf material using a modified CTAB extraction protocol (Doyle and Doyle 1987; Folk and Freudenstein 2014). Standard Illumina TruSeq libraries were constructed at RAPiD Genomics (Gainesville, FL, USA), and target sequences were captured in-house with a 277-locus biotinylated RNA baitset described previously (Folk et al. 2015) and synthesized by Arbor Biosciences. Capture conditions followed the version 4 MyBaits protocol but with modifications identical to those described previously (Folk et al. 2015). Sequencing was performed by RAPiD Genomics, generating 150-bp paired-end Illumina data.

### Assembly

We used aTRAM 2 (Allen et al. 2018) for targeted assembly of nuclear loci. aTRAM is an iterative assembly method that implements a suite of *de novo* assemblers and uses an iterative BLAST process to grow a target sequence from a reference. The assembler used here was SPAdes (Bankevich et al. 2012), with 5 assembly iterations. References for assembly were the original sequences used to design probes. Because probes targeted continuous DNA regions across exons and introns (Folk et al. 2015), most assemblies covered at least the full length of the reference. To identify paralogs, we used a sequence similarity criterion (reviewed in Altenhoff et al. 2019) where the contig with the highest bitscore against the reference (which was within the ingroup) was chosen as the putative ortholog for downstream alignment. This approach represents a compromise in orthology assessment, as distance-based methods are more scalable with taxa than tree-based methods (e.g., Yang and Smith 2014), but they do not explicitly assess homology in a phylogenetic context and may particularly have more difficulty distinguishing in-paralogs (sensu Altenhoff et al. 2019). Phasing has been of particular interest in the phylogenomics community, particularly in polyploids where they could mislead inference (Eriksson et al. 2018; Nauheimer et al. 2021; Karbstein et al. 2022 but see Kates et al. 2018). Because the assembly approach used here is based on a *de novo* algorithm, the resultant data are effectively “phased” (in that alleles, if they differ in sequence, are resolved as separate contigs that are not combined) with only one allele arbitrarily chosen for downstream analysis based on distance. While this approach does not make full use of sequenced allelic data, it greatly simplifies analysis by not attempting to distinguish alleles from paralogs in the absence of synteny data. Because *Asimitellaria* is a clade of allopolyploid origin and forms a suitable test of the assembly procedure, we investigated the impact on the ultimate phylogenetic findings, observing a topology with identical well-supported clades as those found in a previous study where homeologs were fully resolved using a cloning and iterative partitioning procedure (Okuyama et al. 2012). Manual examination of alignments and gene trees revealed that the choice of homeolog among the A and B subgenomes was inconsistent between genes, but within each gene, the homeolog choice was generally consistent and the gene tree topology within *Asimitellaria* was typically similar to the results of Okuyama et al. (2012).

Off-target capture data are often amenable to high-quality assemblies of organellar genomes (Weitemier et al. 2014), where typically >1% of reads are assignable to the chloroplast genome (Folk et al. 2015). Because chloroplast genomes are effectively haploid and all expected paralogs are nuclear or mitochondrial and therefore much lower in coverage, chloroplast genome assembly is straightforward using read-mapping assembly. BWA (Li and Durbin 2009) was used to align off-target reads against the *Heuchera parviflora* var. *saurensis* chloroplast genome (Folk et al. 2015) to generate near-complete assemblies. To generate consensus sequences for each sample, we called variants on resultant read pileups using the BCFtools and VCFtools suites (Danecek et al. 2011), assuming a haploid genome.

### Alignment and phylogenetics

All alignments (nuclear genes, chloroplast genomes) were performed in MAFFT (Katoh et al. 2009), with a gap opening penalty of 3 but default settings otherwise. To handle sites with substantial missing data in nuclear sequences, mostly corresponding to “ragged” ends of alignments, many of which exceeded the length of the original probe region, we removed all sites with at least 90% missing data. All final tree results were rooted with *Peltoboykinia tellimoides*, the most distant of the three chosen outgroups, following Folk et al. (2019).

For nuclear data, we first inferred a maximum likelihood tree in a concatenated framework using RaxML v. 8.2.12 (Stamatakis 2014). Given that previous phylogenetic analyses of the group have failed to find topological differences among standard coding/non-coding and gene-wise partitioning schemes (Folk et al. 2017), this analysis was unpartitioned to optimize computational times. Rapid bootstraps (option “-f a”; Stamatakis et al. 2008) were also calculated to assess support. Chloroplast phylogenetics followed the concatenated methods described above. Individual gene trees were then inferred in RAxML, using unpartitioned GTR-GAMMA models but otherwise identical to the concatenated tree inference. We inferred coalescent trees using ASTRAL-III (Zhang et al. 2018). Although support in the gene trees was often low towards the tips, many backbone gene tree relationships received moderate to good support and were typically consistent with recent phylogenetic work (Okuyama et al. 2012; Folk et al. 2017, 2018b), and therefore we did not filter clades based on support. Coalescent support was measured with local posterior probability (Sayyari and Mirarab 2016). We primarily report ASTRAL-III results using an allele map to recognized species, but a run without the allele map (which was topologically similar) was used to match taxon sampling in the chloroplast phylogeny for gene tree simulations (see below).

### Characterization of ILS

We sought to verify that previous evidence of chloroplast capture is robust to increased taxon sampling. Given that our target of hybridization inference was the chloroplast, putatively evolving as a single coalescent “gene” (c-gene sensu Doyle 1995), and that there is remarkably little evidence of hybridization from nuclear loci alone (Folk et al. 2017), we implemented gene tree simulations under the multispecies coalescent based on nuclear data alone in order to generate predictions of expected incomplete lineage sorting (ILS), and then used this as a null distribution to examine recovered chloroplast clade probabilities. Gene tree simulations were performed in Dendropy (https://github.com/ryanafolk/simulate_gene_trees; based on Mirarab et al. 2014) using the ASTRAL tree without an allele map. For bisexual plants, a coalescent branch length scaling factor of 2 is commonly used to account for expected chloroplast genome *N_e_* (given matrilineal inheritance but both parents yielding offspring; Joly 2012), but given the sporadic occurrence of dioecy and gynodioecy in this group (in *Asimitellaria* and *Tellima*; Folk et al. 2021) and its uncertain implications for ancestral *N_e_*, we also tested a factor of 4 (given both matrilineal inheritance and only female plants yielding offspring, arbitrarily assuming equal sex frequencies; see also García et al. 2017). Finally, we mapped simulated clade frequencies on the empirical chloroplast-based trees.

Chloroplast-based clades that are an expected outcome in the presence of ILS should have non-zero probabilities, while a topology that is poorly predicted by ILS alone (consistent with the presence of hybridization) should have many clade frequencies of probability ∼0. This observation suggests an obvious statistical test for clade frequencies near zero (Folk et al. 2017; García et al. 2017). But even in the case of gene trees generated under ILS, many clades could also have low probabilities when gene conflict is high and many taxa are sampled. To develop an appropriate null distribution representing our clade frequency expectations, we used two approaches. First, we calculated clade probabilities on the simulated gene tree set to characterize the null clade probability expectation, and compared this distribution to the observed distribution of chloroplast-based clade probabilities. Second, we calculated the set of pairwise Robinson-Foulds distances between all pairs of simulated gene trees to characterize the null expectation for the amount of discord and compared this to the empirical distances between the observed chloroplast tree and all simulated gene trees.

### Phylogenetic dating

Given the relatively large number of taxa and genes, we chose MCMC_TREE_ in PAML 4.9 (Yang 2007) to generate a dated phylogeny using nuclear loci. MCMC_TREE_ represents a suitable compromise for phylogenomic data (dos Reis and Yang 2019) between highly parametric methods such as BEAST (Drummond and Rambaut 2007) that would be challenging to run on our dataset without strong reduction of loci and individuals, and fast but relatively simplistic rate smoothing approaches (Smith and O’Meara 2012). We ran MCMC_TREE_ in two analysis setups, with topologies based on either the concatenated or ASTRAL analysis but using identical parameters and nucleotide alignments. For the concatenation analysis, the tree was randomly pruned to one accession per species. Given the absence of a usable fossil record within Saxifragaceae (but a strong record of fossils in close relatives of the family; Magallón et al. 2015), three secondary calibration points were used for time calibration following Deng et al. (2015): the mean ages for the Darmereae [6.68 - 15.53 MYA (millions of years ago)] and Heuchereae [4.43 - 10.56 MYA] clades, and that for the MRCA of Darmereae + Heuchereae + Micrantheae [21.74 - 36.51 MYA]. The probabilities of exceeding calibration upper and lower bounds were set as 0.01. BASEML was used to obtain branch lengths with the GTR+G model. Rate priors on internal nodes were set using autocorrelated rate models. We ran MCMC_TREE_ for 50 million generations as an initial burn-in, followed by 50 million generations, sampling every 1,000 generations. To check for convergence, MCMC_TREE_ was run with random seeds four times for each analysis setup.

### Occurrence records, niche modeling, and climatic data extraction

Occurrences and models used here are from previous occurrence aggregation efforts (Folk et al. 2018b, 2019, 2021). Species were checked again for any potential spatial errors in point records against the taxonomic literature.

Environmental predictors followed those used previously on this dataset (Folk et al. 2018b, 2019) and comprise a broad swathe of 12 predictors relevant to plant distributions. Climate was represented by four variables from Bioclim v. 1 (Hijmans et al. 2005), representing averages of climate data between 1950 and 2000. Monthly data are available through the updated Bioclim 2 product (Fick and Hijmans 2017), but this was deemed unnecessary for the present study, and Bioclim 1 is consistent with previous work (Folk et al. 2018b). Two Bioclim variables represented absolute temperature and its seasonality (Bio1 and Bio7; that is, mean annual temperature and temperature annual range), and two variables represented temperature and precipitation during the dry season (Bio12 and Bio17; that is, annual precipitation and precipitation of the driest quarter). Two topographic variables (elevation and slope; https://doi.org/10.5066/F7DF6PQS), four soil variables (mean coarse fragment percentage, mean pH, mean sand percentage, mean organic carbon content; Hengl et al. 2017), and two variables representing land cover (needle-leaf and herbaceous land cover; Tuanmu and Jetz 2014) were also used to represent other important factors constraining plant distributions.

For taxa with sufficient data (which we considered to be at least 15 vetted occurrence records, chosen such that the number of predictors never exceeded the number of data points), species distribution models were built using Maxent (Phillips et al. 2017), following the parameters in Folk et al. (2018b). Briefly, this began with defining accessible areas for the model training regions using the intersection of a convex hull and occupied ecoregions. Models were trained at full 30-second resolution using 75% of data points, with 25% set aside for model test statistics; this was performed in 10 bootstrap replicates per model with resultant outputs averaged. Extrapolations were disallowed and missing data were allowed; settings otherwise were defaults. Climatic data extraction for ancestral niche reconstruction was performed on averaged models using custom code (https://github.com/ryanafolk/pno_calc). For those taxa with fewer records, given the importance of comprehensive species sampling, we instead extracted climatic data by intersecting occurrence records directly with environmental data. Climatic data extractions, either from species distribution models or from point extraction, were represented as histograms (“predicted niche occupancy profiles” or PNOs, as described previously; Evans et al. 2009).

### Paleoclimate layers

Historical layers for mean annual temperature (Bio1) were reconstructed for 51 time points between and inclusive of 0 and 3.3 MYA using custom scripts to extend a previous high-resolution dataset at selected time points on this interval (Brown et al. 2018). This reconstruction is based on global historical temperature curves with high-temporal resolution (Hansen et al. 2013) to represent equally spaced time intervals covering the Plio-Pleistocene boundary to the end of the Holocene (3.3 MYA to pre-industrial present). Environmental data layers were generated based on five of the layers available through the PaleoClim dataset (Brown et al. 2018) for mean annual temperature at 10 arc-minutes resolution (∼20 km at the equator). These layers were chosen to cover major time points from the Pliocene to the present: specifically, 0 MYA (present), 0.021 MYA (Last Glacial Maximum), 0.787 MYA (MIS19 in the mid-Pleistocene), 3.264 MYA (the mid-Pliocene Warm Period), and 3.3 MYA (Pliocene Glacial Event M2). We conservatively only reconstructed mean annual temperature because historical trends in temperature and precipitation are temporally and spatially distinct but only temperature data were available for extrapolation. This reconstruction method is an extension of the approach of Gamisch (2019), with two primary improvements. First, here we utilize a stronger set of historical time periods through PaleoClim layers, whereas Gamisch (2019) used data from two time points, namely modern and Last Glacial Maximum (∼22 kya). Second, we avoided climatic extrapolation in the prediction, which could be associated with poor predictive performance, by only inferring time points between the PaleoClim layers.

Prior to reconstructing time points, each input layer was cropped to the study area extent and aggregated by a factor of two to reduce spatial resolution and decrease computational times. Universal kriging was then used to calculate geographically interpolated surfaces with spatial linear dependence, especially useful for predicting environmental conditions for areas not included in the original layers (e.g., a newly available coastal region). For each input layer, we determined a variogram where the layer values were linearly dependent on the spatial coordinates (i.e., in R notation, bio ∼ longitude + latitude) with the autofitVariogram function from the R package automaps (Hiemstra, P., and M. P. Hiemstra 2013). Next, universal kriging was implemented using the krige function from the R package gstat (Pebesma 2019).

Layers were then inferred for the indicated time periods as follows. First, we identified two input layers with the most similar surface temperature associated with the desired time point through linear interpolation on the temperature curve produced by Hansen et al. (2013) with the R function approx. We then used these layers to calculate a Δ layer between the identified input layers using the overlay function from the R package raster (Hijmans and van Etten 2012). Specifically, the Δ layer is equal to the difference between the layer with the closest minimum surface temperature and the layer with the closest maximum surface temperature.

We then calculated and applied a surface temperature correction. The surface temperature correction was calculated based on surface temperature values identified with linear interpolation on the temperature curve produced by Hansen (2013). We identified the approximate surface temperature value for the time being reconstructed or the temperature surface initial (T_SI_), as well as the surface temperatures associated with the two input layers with the most similar surface temperature, the closest minimum surface temperature (T_SA_), and the closest maximum surface temperature (T_SB_). The surface temperature correction is equal to (T_SI_ - T_SB_)/(T_SA_ - T_SB_). The Δ layer was then multiplied by the surface temperature correction, and the resulting layer is referred to as the ΔT layer. The layer was then corrected with the Delta method (Ramirez-Villegas and Jarvis 2010), which is often applied when downscaling climate layers; here, the layer of the closest maximum surface temperature (T_SB_) was added to the ΔT layer.

Using the ETOPO1 Global relief model (Amante and Eakins 2009), we corrected coastlines for each time period examined. ETOPO1 extent was cropped to match the Bio layers (180° E, 180° W, 0° N, and 90° N). Additionally, the resolution of the ETOPO1 layer was set to match the Bio layers using a nearest neighbor approach with the projectRaster function in the R package raster. We corrected the raster values for ETOPO1 by adding an approximated sea level change as provided by Hansen et al. (2013). We reclassified anything between 0 m and −15,000 m as water (i.e., as 0) and any value above 0 m as land (i.e., as 1; Hansen et al. 2013; Gamisch 2019). This raster was then multiplied with the scaled ΔT layer to correct the coastline, which utilizes the modified ETOPO1 layer as a mask.

The performance of paleoclimatic interpolation was assessed via a jackknife approach, successively removing one layer at a time among the input layers and predicting the missing raster. We then verified high overall similarity between the predicted and PaleoClim layers based on recovering a Pearson correlation coefficient above 0.70.

### Ancestral reconstruction of environmental tolerances

We implemented several new methods for generating predictions of potential distributions based on climatic data. Many approaches exist (e.g., Yesson and Culham 2006; Evans et al. 2009; Folk et al. 2018b) for so-called “phyloclimatic modeling,” which generally comprises inference of climatic occupancy in present-day species, integrating this in various ways with ancestral reconstruction methods, and projecting reconstructed climatic occupancy onto paleoclimatic reconstructions. One of the key differences among these approaches lies in how they handle the niche breadth of extant species. While accounting for trait variation is a common problem, and often no more than a biological annoyance, in ancestral reconstruction and other comparative methods (Wiens 1999; Felsenstein 2008), species never occupy a single point in ecological space, and therefore niche reconstruction is generally incomplete under point-based methods (Saupe et al. 2018). Variation in the occupancy of niche space, referred to in this context as niche breadth, is intellectually central to the concept of niche and possibly more interesting than the niche estimate itself depending on the research questions (Sexton et al. 2017). Some representative examples of methodological approaches to estimating niche breadth attempted so far are min-max coding (Yesson and Culham 2006) and bootstrapping over environmental data samples (Evans et al. 2009; Folk et al. 2018b). Min-max coding has important limitations such as the questionable homology of the extreme limits of a multidimensional trait, the sensitivity of these statistics to incomplete modeling of the niche (Saupe et al. 2018), and the failure to incorporate the distributional shape of environmental occupancy. Environmental occupancy breadth could be defined via bootstrapping approaches, but in practice bootstrapping does not meaningfully reconstruct the shape of this distribution because the resultant ancestral reconstructions tend to be normally distributed (e.g., see Folk et al. 2018b: Fig. 3).

We developed a new ancestral reconstruction approach based on histogram statistics calculated on predicted niche occupancy profiles (that is, probability density profiles that represent niche as a response to a single environmental variable, sensu Evans et al. 2009). Instead of drawing from the distribution of environmental data for many independent reconstructions, we divide the environment into a series of bins shared across species and reconstruct the height of these bins across species (Fig. 1a). The reconstruction method is from this point forward a standard implementation of Brownian motion under maximum likelihood. This treatment carries the assumption that each bin evolves as an independent character that is allowed to have independent rate parameters and root states. The assumption of independence is a strong one, but is similar to assumptions of site independence in standard evolutionary models of multiple sequence alignments and reasonable, particularly in the motivating case of complex niche distributions. An ancestral reconstruction philosophy that more directly represents the shape and breadth of extant niche allows the possibility of allowing asymmetry and multimodality in the niche of present-day species to inform ancestral reconstruction. In practice, the histogram approach enables a natural way to reconstruct arbitrary distributional shapes of environmental tolerance in ancestral taxa. Among previous approaches, that of Saupe et al. (2018) is most similar, but it represents a presence-absence approach and does not attempt to integrate over the probability of presence represented in PNOs. We added this code, implemented in Python, to the BiotaPhy Analyses repository (https://github.com/biotaphy/BiotaPhyPy/blob/main/biotaphy/tools/ancestral_distribution.py; original version by S.A.S. and B.C.O. at https://github.com/blackrim/anc_distr_rec) so that future users can utilize the method through the BiotaPhy platform (https://data.lifemapper.org/biotaphy/; Soltis and Soltis 2016) or locally on their own data.

**Fig. 1.**
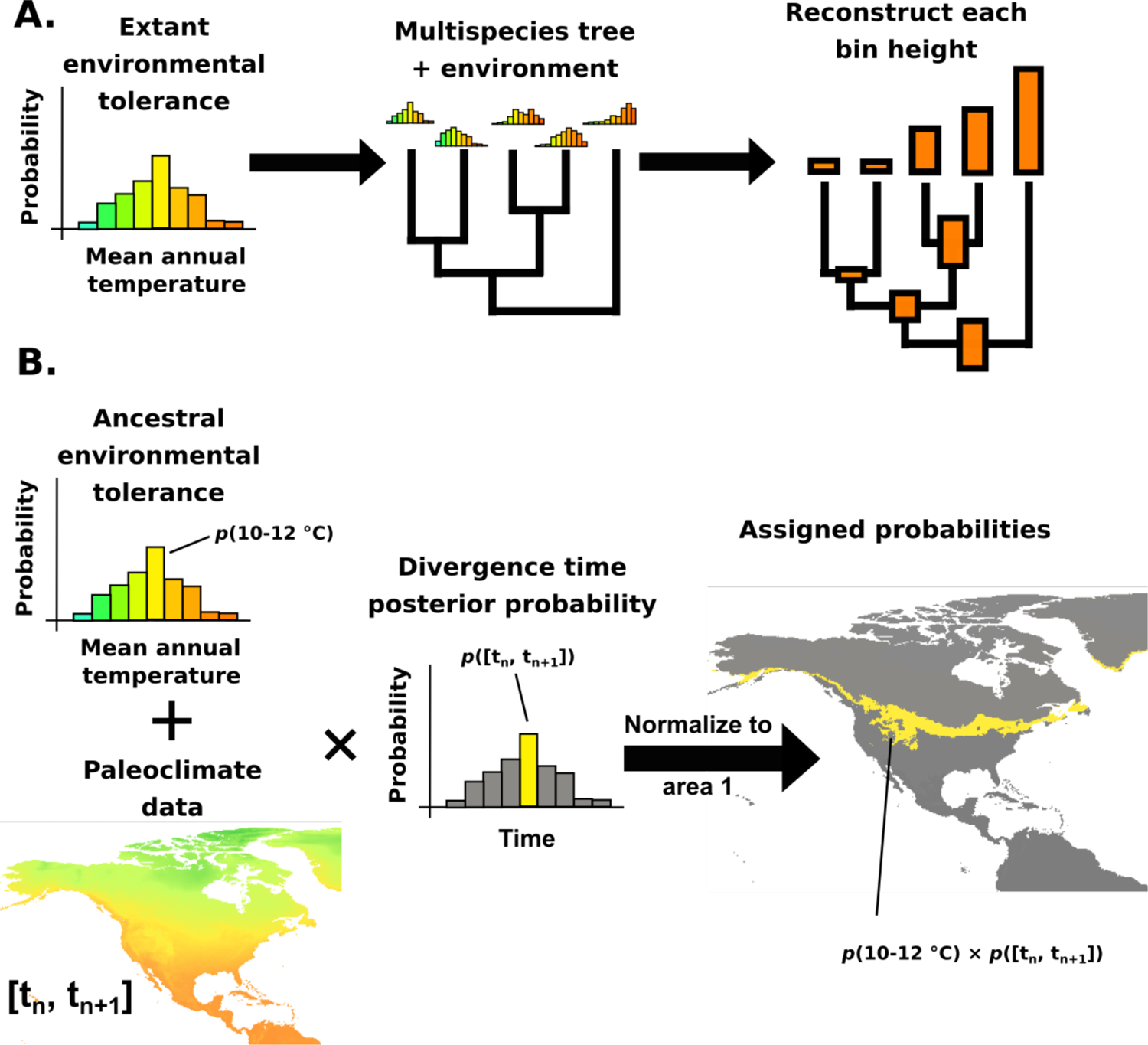
Workflow for ancestral niche reconstruction. (A) Steps for the binned ancestral reconstruction for a single bin. (B) Steps for geographic projection of the binned ancestral reconstruction result for a single node, where *t* represents divergence time.

### Paleo-range prediction

With reconstructions of past environmental tolerance in hand, we generated potential past range predictions using the interpolated paleoclimatic layers discussed above. We chose to focus paleoclimatic data on mean annual temperature, an important predictor of plant distributions as well as the predictor with the best performance among our paleoclimatic interpolations based on the jackknife procedure (see above), so these predictions represent the dynamics of broad bands of temperature tolerance since the mid-Pliocene. Notably, our reconstruction of mean annual temperature is still conditional on the full variable set used in the niche model predictor set, with the simplifying, albeit imperfect, assumption that this conditional relationship among predictor variables is constant through time.

Several methods have been published for translating environmental tolerance reconstructions into past range predictions, generally binary predictions based on min-max coding (Yesson and Culham 2006), distance methods (Meseguer et al. 2018; Williams et al. 2018), or methods based on histogram statistics (Folk et al. 2018b); the last approach has the advantage of being directly interpretable as a habitat suitability metric (unlike unbounded distance metrics) that represents probability at relatively fine grain (unlike coarse binary maps). Methods for projection here use histogram statistics following Folk et al. (2018b) and naturally complementing the histogram method for ancestral reconstruction. Briefly, the paleoclimate raster was paired with corresponding bins in the PNO histograms and then classified by the corresponding bin probabilities (Fig. 1b). This was done on a per-species basis, and predicted ranges were summed across species. Finally, while hybridization has also been documented for East Asian species (Okuyama et al. 2005; 2012), to capture the research scope of this contribution (that is, evaluating the hypotheses of Folk et al. 2017; 2018b) the prediction was trimmed to North America, and the histogram was normalized to area 1 to yield probability densities.

An important problem arises in linking climatic data with phylogenetic time calibration: uncertainty in phylogenetic dating tends to be very large compared to the relatively narrow temporal scales represented by recent Pleistocene climatic shifts (reviewed in Folk et al. 2018b; see especially Fig. 6 in that paper). For instance, glacial-interglacial periods are measured in tens of thousands of years, but a dating uncertainty of 1 million years or more is not uncommon even in fairly recent divergences. It is not ideal to use point estimates under these conditions. We addressed this mismatch in temporal scale by explicitly integrating dating uncertainty into our paleoclimatic projections, implemented as another set of histogram statistics. We used Dendropy (Sukumaran and Holder 2010) to derive histograms representing posterior date probabilities from MCMC_TREE_, fixing the bin boundaries to correspond to the 51 developed for paleoclimatic layers (discussed above). We used these bin probabilities to weight paleoclimatic predictions for every node in the phylogeny by the posterior probability of occurring in each of the 50 time frames and projected all nodes into all time frames. Then for each time frame, the sum was taken for all nodes (analogous to species richness maps, but here summing occurrence probabilities instead of species counts). Nodes certain not to occur in a given time frame have posterior probability ∼0 and drop out, while nodes possibly occurring in multiple time frames are projected across them proportional to probability to adequately represent uncertainty. We did not incorporate phylogenetic uncertainty; we feel this is justified given the small number of poorly supported clades in the topology obtained in this study. Notably, our results (below) are generally comparable to a previous investigation that did integrate phylogenetic uncertainty using Bayesian methods (Folk et al. 2018b). Nevertheless, the method could be extended to incorporate uncertainty by averaging the results over a collection of trees representing uncertainty, such as bootstrap tree samples, to obtain predictions weighted by clade uncertainty.

### Comparison with published methods

We compared the centroid and niche breadth estimates of the histogram method proposed here with two previously published methods: (1) a widely used ML implementation in R package Phyloclim (Heibl 2011) and (2) a recent Bayesian implementation in Ambitus (Folk et al. 2018b). These packages were run using an identical tree and PNO data with default settings to represent typical usage.

### Data availability

Scripts for performing ancestral reconstruction analyses, other than those given above, are posted on GitHub (https://github.com/ryanafolk/heuchera_ancestral_niche/; https://github.com/ryanafolk/simulate_gene_trees). The described approach for interpolating paleoclimate layers is also posted on GitHub (https://github.com/mgaynor1/PaleoGenerate), using only open access datasets as specified above. Phylogenies, gene trees, alignments, and paleoclimatic interpolations are published at Dryad (XXX). Raw sequence data are published on SRA (PRJNA641968). Note to reviewers: Animated GIFs representing trends in suitable habitat over time are available at https://github.com/ryanafolk/heuchera_ancestral_niche/blob/master/results/ancestral_projection_animation_ASTRAL_tree/combined_constantscale.gif (ASTRAL tree) and https://github.com/ryanafolk/heuchera_ancestral_niche/blob/master/results/ancestral_projection_animation_concatenation_tree/combined_constantscale.gif (concatenated tree); we ultimately aim to distribute these as social media materials and as embedded multimedia materials in the online version of this manuscript.

## Results

### Nuclear-based phylogenetics

Clade support values were generally high (BS [bootstrap] ∼100; LPP [local posterior probability] ∼ 1), especially in concatenation analyses. Members of Heuchereae, itself confidently resolved as monophyletic, were resolved in one of three major clades, namely *Heuchera,* the “*Ozomelis* group,” and the “*Pectiantia* group” (Figs. 2-3; the last clade interpreted differently from Folk et al. 2014). Other than population samples of the same species, most phylogenetic uncertainty in *Heuchera,* as measured by support estimates and observed incongruence between analyses, was restricted to close relatives in *Heuchera* section *Rhodoheuchera*, as observed previously (Folk et al. 2017). Most relationships were consistent with previous work; differences from previously published studies are discussed briefly here, and overall phylogenetic results and comparisons with previous studies are reported in more detail in Supplemental Table S4. Within *Heuchera* sect. *Rhodoheuchera*, we recovered two novel relationships that were restricted to concatenation analysis (Fig. 2) and disagreed with previously published work including concatenation approaches. First, *Heuchera bracteata* and *H. hallii* appear embedded within *Heuchera* sect. *Rhodoheuchera* rather than as a clade sister to *H. woodsiaphila* + *Heuchera* sect. *Rhodoheuchera* (as seen in the coalescent analysis [Fig. 3] and in Folk et al. 2017); all branches resolving this relationship have only moderate bootstrap support (<75%). Second, *Heuchera caespitosa,* resolved as embedded within *Heuchera* subsect. *Elegantes* in the coalescent analysis and in previous work (Folk et al. 2017, 2018b), had a different position as sister to *H. rubescens* var. *alpicola*, with these two taxa forming a clade sister and reciprocally monophyletic with respect to *H. bracteata* and *H. hallii*.

**Fig. 2.**
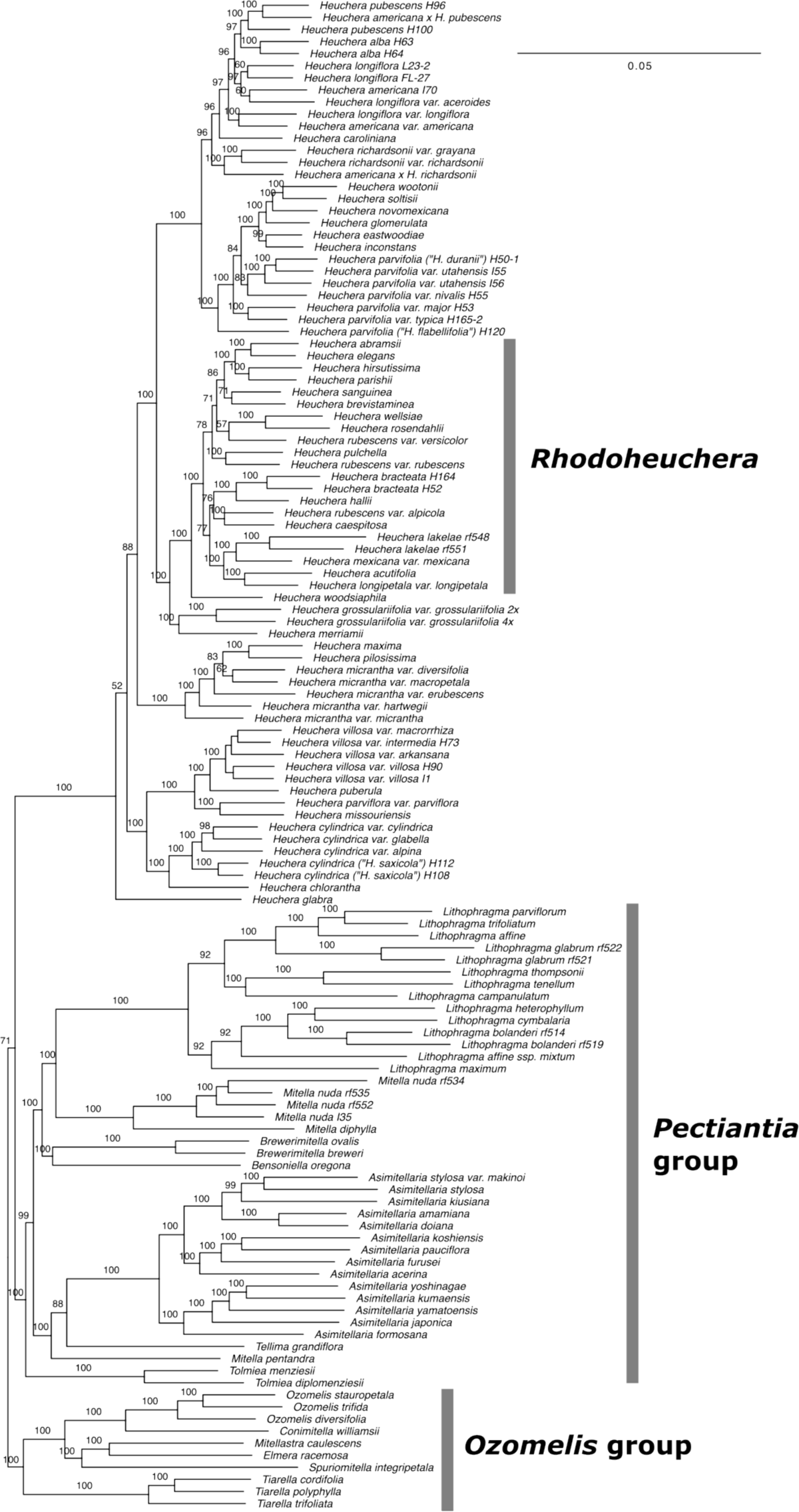
Topology recovered under concatenation in RAxML using the 277 nuclear loci. Branch labels represent bootstrap percentages ≥ 50%. Branch lengths represent per-site substitutions. Informal clade names noted in the text are labeled with gray boxes. Outgroup branches (*Darmera, Rodgersia, Peltoboykinia*) are omitted for compact representation.

**Fig. 3.**
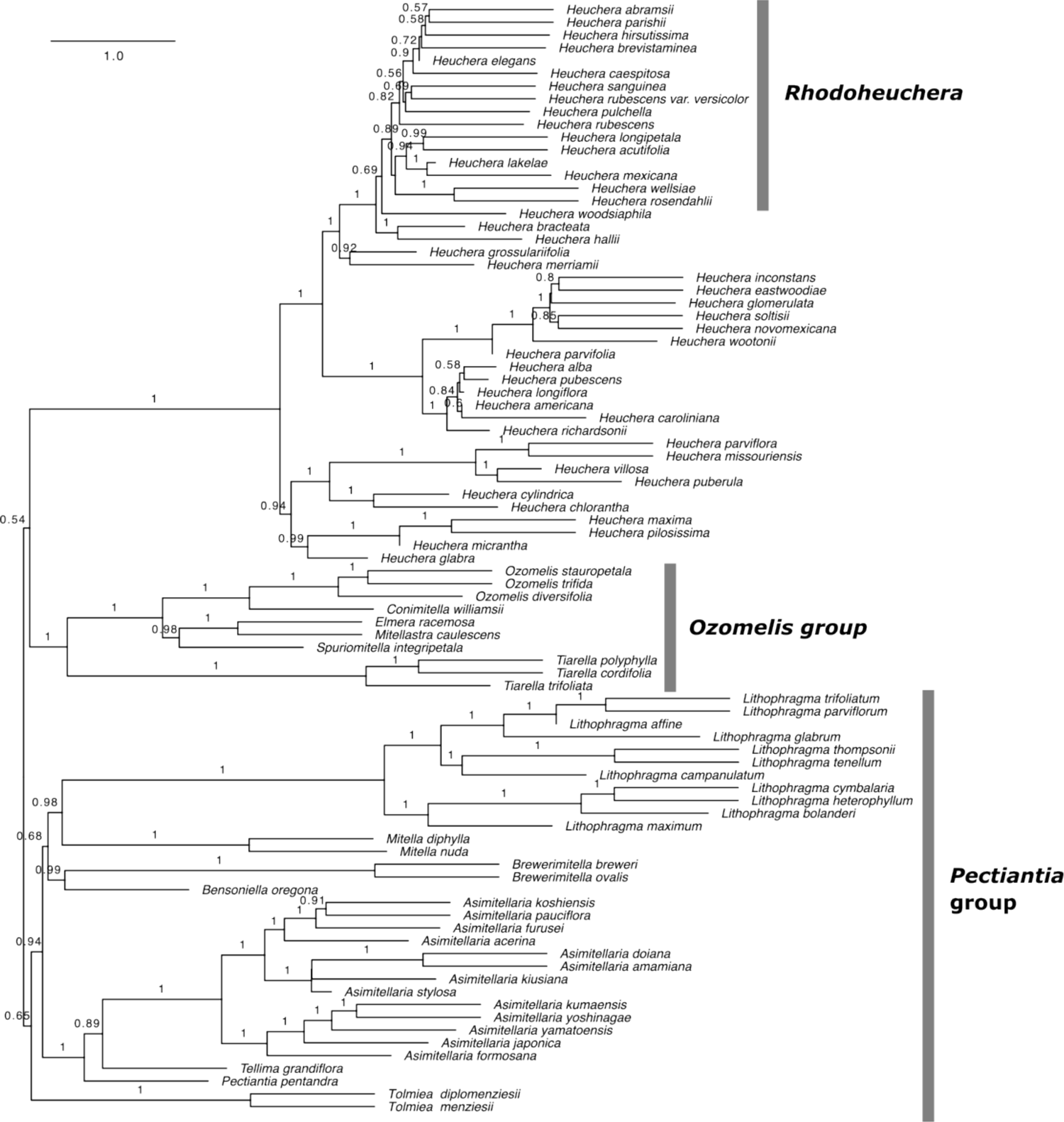
Topology recovered under coalescence in ASTRAL using the 277 nuclear loci. Branch labels represent local posterior probabilities (Sayyari and Mirarab 2016) and were only plotted for those nodes with ≥ 0.5 probability. Branch lengths represent coalescent units; tip taxa without sampling of multiple individuals are arbitrarily plotted as 1.0. Informal clade names noted in the text are labeled with gray boxes. Outgroup branches (*Darmera, Rodgersia, Peltoboykinia*) are omitted for compact representation.

Two further results disagreed between concatenation and coalescent analyses but were already known to disagree among previous studies. The *Ozomelis* group, although only recovered with moderate support in our concatenated analysis (BS 71), was sister to all other species of tribe Heuchereae, congruent with results recovered previously using ribosomal ITS and ETS loci (Okuyama et al. 2008; Folk and Freudenstein 2014). Coalescent analyses instead recovered *Heuchera* as sister to the *Ozomelis* group with weak support (LPP 0.54) as seen in one of the analyses of Okuyama et al. (2012). Finally, *Heuchera glabra* was sister to the rest of the genus in the concatenation analysis, as seen in a previous total evidence analysis including morphology but not in DNA-only analyses (Folk and Freudenstein 2014), whereas in the coalescent analysis it was sister to a clade containing *H. maxima*, *H. micrantha*, and *H. pilosissima* (that is, *Heuchera* subsect. *Micranthae* sensu Folk 2015), in agreement with previous phylogenomic studies that included this species (Folk et al. 2017, 2018b). Otherwise, relationships among and within genera other than *Heuchera* were highly congruent and largely well-supported across all methods. Several further relationships in the ASTRAL tree that did not receive high local posterior support were congruent with decisively supported nodes in the concatenation tree. Among taxa not included in previous nuclear-based phylogenomic studies, *Conimitella, Spuriomitella,* and additional species of *Asimitellaria*, *Heuchera, Lithophragma, Ozomelis,* and *Tiarella* were placed as expected based on previous studies (Okuyama et al. 2012; Kuzoff et al. 1999; Liu et al. 2020).

Chloroplast DNA results, with increased sampling relative to previous studies, were essentially identical in topology to previous phylogenomic investigations of the chloroplast genome (Folk et al. 2017, 2018b; Liu et al. 2020), to previous results based on the *trnL-F, rpl32-trnL*, and *rps16-trnK* regions of the chloroplast genome (Folk et al. 2017), and, aside from backbone relationships, to results based on restriction site variation (Soltis et al. 1991). Chloroplast DNA relationships differ greatly from the nuclear-based phylogeny, as described at length previously (Soltis and Kuzoff 1995; Folk et al. 2017). Nuclear- and chloroplast-based trees both resolve the monophyly of tribe Heuchereae and the relatively small genera *Asimitellaria, Brewerimitella, Lithophragma, Mitella,* and *Tolmiea.* Yet backbone relationships among these genera are completely different and strongly supported in the chloroplast- and nuclear-based trees (see chloroplast DNA analysis support values in Supplemental Fig. S1), with numerous well-supported topological differences within genera (compare Figs. 2 and 3 to Fig. 4). Notably, species of *Heuchera*, recovered as confidently monophyletic in every major nuclear phylogenetic study of the genus conducted to date, appear dispersed among three distantly related and well-supported major clades in chloroplast-based trees. These clades, named A, B, and C in Folk et al. (2017), were all recovered here with decisive support. The increased taxon sampling of this study placed the monotypic genus *Spuriomitella* as sister to also monotypic *Tellima* (not in the *Ozomelis* group as in the nuclear-based analyses); additional species of *Lithophragma* and *Asimitellaria* were placed close to previously sampled species (similar to nuclear analyses), and *Heuchera lakelae*, not sampled previously, was placed in chloroplast clade B close to other Mexican species (similar to nuclear analyses). A novel topological result placed a monophyletic *Tolmiea* within *Lithophragma*, sister to a clade comprising *L. bolanderi, L. cymbalaria,* and *L. heterophyllum*.

**Fig. 4.**
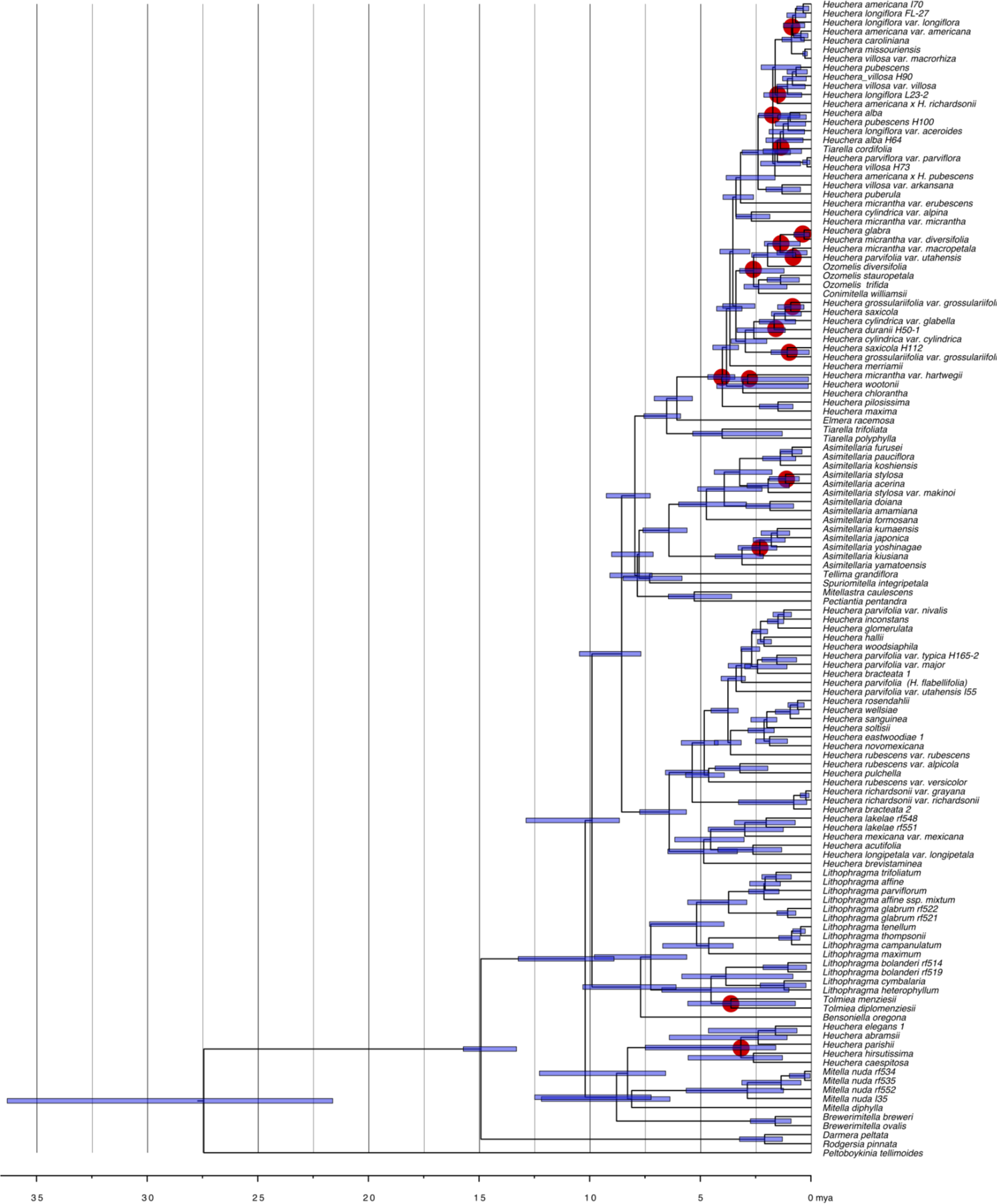
Dating result for the chloroplast phylogeny from MCMC_TREE_ using the 277 nuclear loci. The x-axis represents time in millions of years; node bars represent 95% credibility intervals. Red circles denote MRCA dates for chloroplast capture events inferred in Folk et al. (2017), as well as the event inferred here for *Tolmiea* (see Results). Outgroup taxa are included here (cf. Figs. 2, 3).

Dating results for nuclear data were similar among the four replicates of each dating analysis, and also between concatenation and coalescence analysis given that the backbone topology was nearly identical. To highlight major recognized clades, the analysis of the ASTRAL topology (Fig. S2) yielded 11.51 MYA, with 95% credibility interval [11.30, 11.98 MYA] for the MRCA of tribe Heuchereae (i.e., the taxa given in Figs. 2 and 3), 9.50 MYA [8.87, 10.00 MYA] for *Heuchera*, 10.85 MYA [10.57, 11.19 MYA] for the *Ozomelis* group, and 10.34 MYA [10.07, 10.61 MYA] for the *Pectiantia* group. While we ran downstream ancestral reconstruction analyses on both nuclear trees, for the purpose of discussion, we focus hereafter on coalescence results for the nuclear genome unless otherwise noted. The chloroplast MRCA date for tribe Heuchereae, which was among the dating constraints, was 10.20 MYA [8.90, 13.23 MYA]. Chloroplast data tend to recover younger divergence dates than nuclear data in this group (Folk et al. 2018b); dates for the unconstrained clades recovered by both genomes were mostly similar and are fully reported in Table S2.

### Coalescent simulation

We used a coalescent simulation approach based on ILS expectations to test whether increased sampling supported the hypothesis of chloroplast capture as an explanation for cytonuclear discord. Consistent with previous work that had limited taxon sampling outside of *Heuchera* (Folk et al. 2017), we find that the chloroplast topology recovered here is extremely unlikely given the nuclear gene tree distribution and ILS alone, with all backbone clades having probability ∼0 in chloroplast gene tree simulations. The empirical distribution of clade probabilities in the chloroplast tree was significantly different from the null expectation (one-tailed *t*-test with equal variance; *p* = 6.77e-5). Measured as Robinson-Foulds distances, the comparison is also significant (one-tailed *t*-test with equal variance; *p* < 1e-20). As mentioned above, chloroplast-nuclear phylogenetic discord was very similar to that found previously (Folk et al. 2017, 2018b), and overall simulation results were also similar, but our complete sampling of *Lithophragma* enabled the detection of a new putative chloroplast capture event, given that *Tolmiea* is nested within *Lithophragma* (Fig. S1). Both genera are morphologically well defined, and the latter has long been recovered as monophyletic and not closely related to *Tolmiea* with nuclear markers (Kuzoff et al. 1999; this work). This new topological placement is also unexpected under ILS alone (chloroplast clade frequency < 10e-9; best alternative relationship 0.047). We also corroborated the hypothesis of Okuyama et al. (2005) of two chloroplast capture events within *Asimitellaria* (Fig. 4).

### Ancestral niche projection

The reconstructed distribution of suitable habitat for Heuchereae shifted southwards in cool periods of the Pleistocene, and predictions of range restriction primarily occurred during the mid- to late Pleistocene. The first southward refugial distribution was reconstructed at approximately 1.65 MYA (Fig. 5; see also https://github.com/ryanafolk/heuchera_ancestral_niche/blob/master/results/ancestral_projection_animation_ASTRAL_tree/combined_constantscale.gif), followed by repeated oscillations between broad and restricted predicted species ranges to the present. An important limitation of translating ancestral niche reconstructions into species distributions is the often high uncertainty associated with time calibration of typical empirical studies. Despite weighting ancestral range predictions by phylogenetic dating uncertainty, we were able to produce high-resolution ancestral range predictions that clearly distinguished among cold and warm periods of the Pleistocene (Fig. 5).

**Fig. 5.**
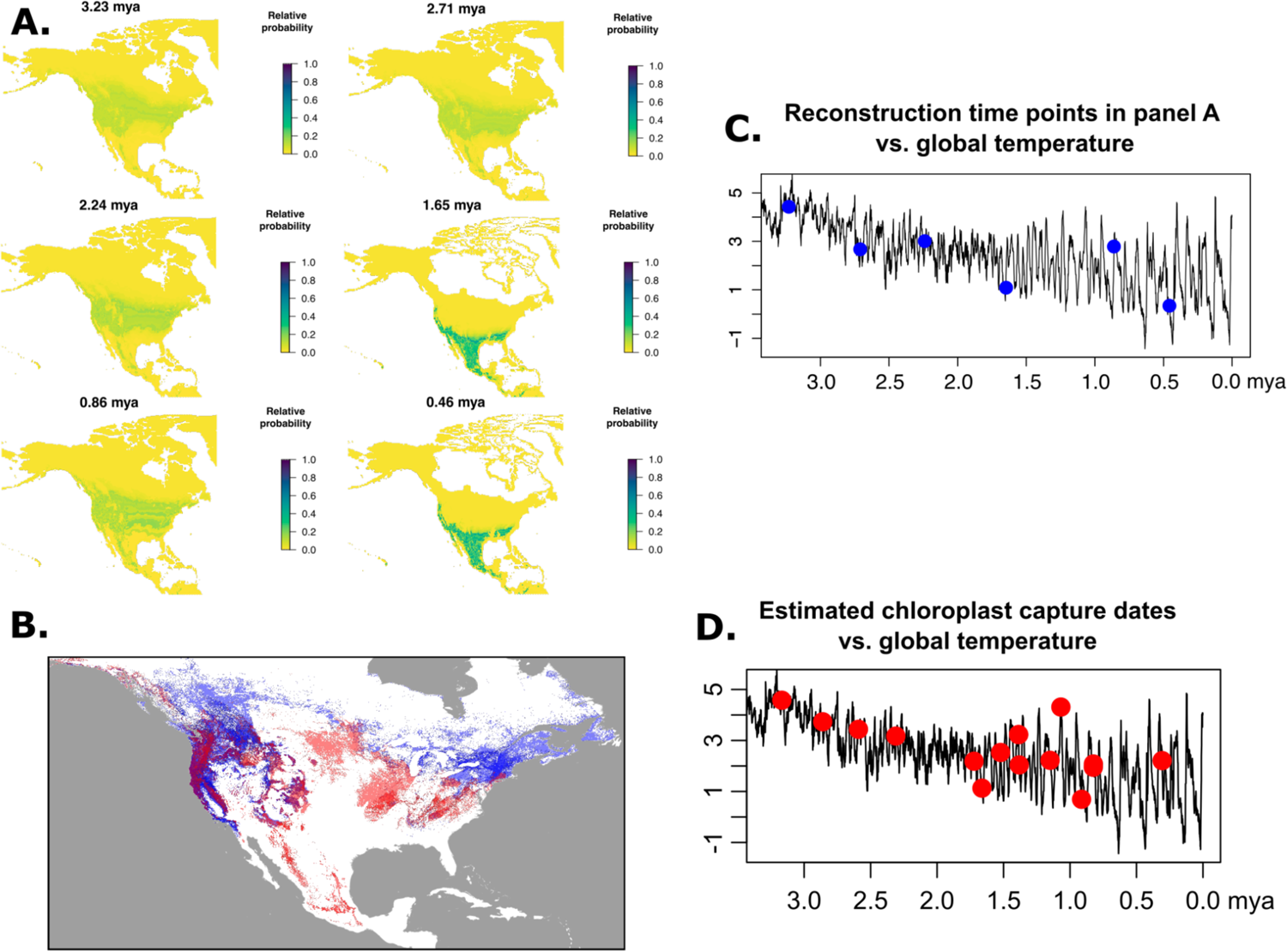
Trends in habitat suitability over time for Heuchereae. (A) Representatives of the 50 ancestral projections between mid-Pliocene conditions (∼3.3 MYA) and the present on the ASTRAL topology. Results are also available as animated GIFs at https://github.com/ryanafolk/heuchera_ancestral_niche/. (B) Present-day distribution of members of Heuchereae estimated by overlain distribution models. *Heuchera* species distributions are shown in transparent red, and other genera of Heuchereae are shown in transparent blue; darker colors indicate greater species richness. (C) Times and global temperatures for each of the six projections in panel (A), plotted on the Zachos et al. (2001) temperature curve; the x-axis represents millions of years to present; the y-axis represents °C. (D) Inferred times of chloroplast capture in the study region, based on reconciling nuclear data with Supplemental Fig. S1, plotted on the Zachos et al. (2001) temperature curve; the x-axis represents millions of years to present; the y-axis represents °C.

Following the chloroplast capture events previously hypothesized (Folk et al. 2017) and confirmed here, the range restriction noted above was concurrent with the date of many previously inferred chloroplast capture events, i.e., primarily within the last 1.75 million years (Folk et al. 2017; plotted in Fig. 5d; see also Fig. 4). To test for a relationship with temperature, we performed a linear regression of paleoclimatic temperature (Zachos et al. 2001) vs. MRCA clade dates for inferred chloroplast capture, finding a significant relationship (*F*-test, *p* = 0.0013; *R*^2^ = 0.4529). A simple observation of correlation between the timeline of hybridization and the Pleistocene could be confounded by the branch length structure of the tree, since branches are denser towards the present; this is a form of autocorrelation that might undermine a direct relationship to temperature. To test whether this distribution of timings was different from the null expectations, we scored a binary matrix of taxa with a history of chloroplast capture events following Fig. 4 and used the R package geiger (Harmon et al. 2008; Pennell et al. 2014) to fit an irreversible discrete model. Based on this rate matrix, we simulated 1000 binary characters and performed stochastic mapping to generate a null distribution with no temporal patterning. The mean MRCA clade age for inferred chloroplast capture events was 1.8166 MYA; this was significantly different from the null expectation (6.8102 MYA, one-tailed equal variance *t*-test, *p* = 0.0055).

## Discussion

### Hybridization and the Pleistocene

The reconstructed dates for many chloroplast capture events in Heuchereae (Fig. 4, enumerated in Table S3) correspond to the Pleistocene epoch, and particularly to after the mid-Pleistocene (within the last 1.75 MYA), a timeframe near the Mid-Pleistocene Transition (1.25 MYA) that is associated with elevated temperature fluctuations and rapid range shift dynamics (Fig. 5; 9 of 12 putative events are within this time frame). The time frame we recovered is therefore consistent with identifying Pleistocene glaciation as a key climatic driver for hybridization opportunities (e.g., Anderson and Stebbins 1954). Our overall time frame is consistent with what has been recovered previously (Deng et al. 2015; Folk et al. 2018b); while Heuchereae lacks a fossil record, as argued by Okuyama (2016), the split between *Asimitellaria amamiana* and *A. doiana* can be constrained as >1.3 MYA based on the geological history of the Ryukyus, which is concordant with the divergence time recovered here.

Within this Pleistocene time frame, ancestral range predictions show a broad contraction and southerly movement of geographic distribution for Heuchereae across its entire range. The geographic distribution of these potential refugial areas is centered on Mexico in an area partly corresponding to its current Mexican distribution, and additional areas that form part of its modern distribution along the coast of western North America and in a disjunct area of the southeastern United States. Distributional contractions corresponded to glacial cycles, with increased range overlap during these times as suggested by occurrence probabilities. Such an association is attributable to novel patterns of range contact spurred by rapid migration during past climate change. Geographic ranges of lowland plants in the Northern Hemisphere typically experienced southward migration and fragmentation of populations in Pleistocene refugia (reviewed in Folk et al. 2018b; see also Soltis et al. 2006; Soltis et al. 1997; Brunsfeld et al. 2002), which would have resulted in potential contact among many species that were previously allopatric but shared refugial areas. But an alternative and important component of a colder Pleistocene climate is the diversity of responses to climate change: while many lowland species are thought to have experienced range restriction, many high alpine taxa had increased habitat suitability and experienced increased range sizes (Guralnick 2006; Folk et al. 2018b). The presence of both ecological strategies in Heuchereae, therefore, would have promoted novel patterns of range contact not evident in present-day populations.

Consistent with a previous small-scale ancestral niche modeling investigation (Folk et al. 2018b) and with verbal models (e.g., López-Alvarez et al. 2015; Klein and Kadereit 2016; Marques et al. 2016), there appears to be a general relationship in *Heuchera* between historical temperature dynamics and opportunities for hybridization facilitated by significant range contractions of many taxa into shared refugial locations. This relationship agrees with the prediction that opportunities for hybridization are facilitated by ecological disturbance (Anderson and Stebbins 1954), a general principle of which Pleistocene climate change is one particularly dramatic example.

### Trends in niche evolution

While most ancestral reconstructions of niche appeared approximately normally distributed, the histogram method was able to construct complex distributions such as bimodality and long tails (Fig. 6c). As expected, uncertainty in trait reconstruction is greatest for deeper nodes. For instance, focusing on temperature, precipitation, and elevation trends in the genus *Heuchera* (results for further genera are available in Table S5), the MRCA of *Heuchera* was reconstructed as living in warm-temperate (maximum bin probability, mean annual temperature 12.0°C, corresponding to, for example, today’s mid-South region of the U.S.A.; see Fig. 6) and wet environments (annual rainfall 1,119 mm, similar to some areas of the present-day South and Midwest regions of the U.S.A.) at low to mid-elevation (this was a multimodal reconstruction with peaks from 84 to 1956 m). Mean annual temperature did not show strong trends as there are shifts to both cooler and warmer environments. Precipitation showed strong trends towards species’ invasion of drier environments across multiple lineages, as did elevation (Fig. S3).

**Fig. 6.**
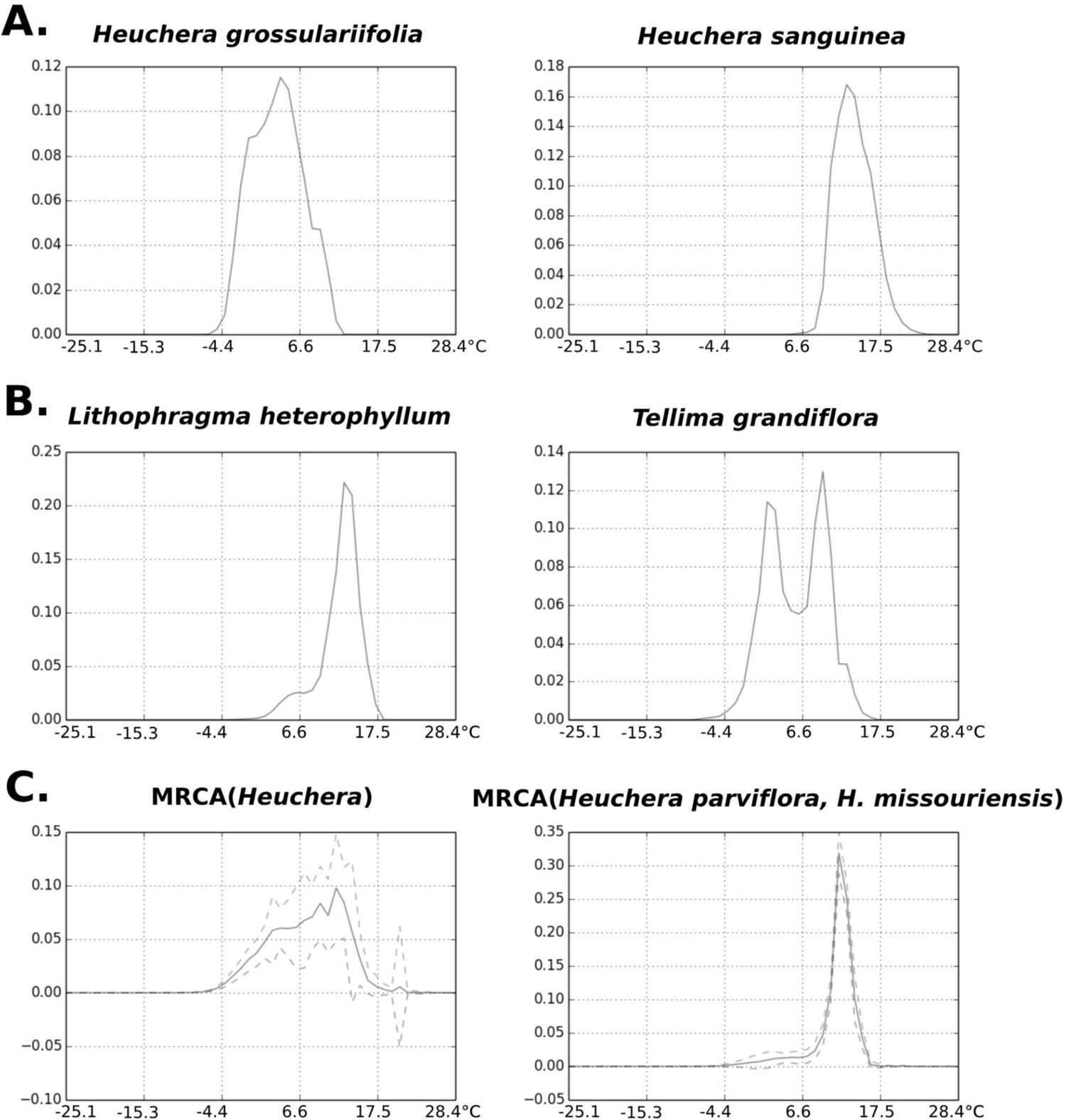
Examples of extant environmental occupancy and reconstructions for exemplary cases on the ASTRAL topology, as histograms here plotted as line graphs for clarity; full plots available at https://github.com/ryanafolk/heuchera_ancestral_niche/. (A) Two examples of near-normal tolerances in extant species. (B) Examples of long tails and bimodality of tolerance in extant species. (C) Examples of ancestral reconstructions of environmental tolerance for a deep node in the tree and a shallow node ancestral to a sister-species pair. Solid lines represent probability densities; dotted lines represent standard errors around these densities. The x-axis in all cases represents mean annual temperature in °C; y-axes represent probability density.

### Histogram methods for ancestral reconstruction

Ancestral niche reconstruction has attracted considerable interest given its importance for understanding past geographic range and its complementarity to standard biogeographic models that typically do not incorporate habitat suitability (Yesson and Culham 2006; Evans et al. 2009; Meseguer et al. 2015; Sukumaran and Knowles 2018; Williams et al. 2018; Landis et al. 2021). The difference between these many approaches primarily centers on the unique “volumetric” aspects of species niche compared to other types of trait data*—*fundamentally they differ in how niche breadth and “shape” (how environmental tolerance varies along the niche breadth) are incorporated. Here we implemented for the first time a histogram approach, where the probability density of predefined environmental bins is the quantity subject to ancestral reconstruction. Such an approach has the advantage of explicitly attempting to reconstruct not only how wide the niche is but its shape, including various forms of non-normality such as multimodal distributions, as are often found in empirical niche models (e.g., see Evans et al. 2009). Traditional approaches such as min-max coding focus on the extremes: the very limit at which individuals of a species survive (perhaps partly due to immigration of those in less extreme habitats). The histogram approach focuses on the whole distribution, de-emphasizing while still including the limits. We find the method can identify potential multimodality in several parts of the tree, particularly in the early history of Heuchereae. This is perhaps consistent with later specialization of the clade into different lithophytic environments across elevational and precipitation gradients, although the histogram method could also disproportionately favor broad ancestral niche predictions (see below). The methods included here will be an important addition to the toolkit for examining habitat evolution in a macroevolutionary context, given the importance of incorporating niche breadth (Saupe et al. 2018).

### Comparison with previous methods

Phyloclim (Heible 2011) does not implement estimates of niche breadth, but the histogram approach developed here results in consistently larger niche breadth estimates than in Ambitus (Folk et al. 2018b; Table S5), reflecting the very different interpretation of niche breadth in the two methods. While this difference represents how a user might interpret the result, it should be noted that the niche breadths reported in Table S5 (range of bins having at least 1% probability and 95% credibility intervals, respectively) represent different calculations and are not necessarily directly comparable. Estimates of the niche centroid, by contrast, were quite similar between methods. The histogram method returned higher estimates overall, but this difference was not significant in comparison with Phyloclim (*p* = 0.1372, paired two-tailed *t-*test, equal variance) and only 0.67 °C greater. Ambitus returned node values averaging 1.2 °C lower than the histogram method, and while modest, this comparison was significant (*p* = 0.0047), likely reflecting the left-skew of most ancestral reconstruction histograms. Overall, while the histogram method differed by some measures reflecting its differing methodological aims, its performance in estimating niche centroids was fairly similar to previously published methods.

### Updated phylogenetic relationships

We have incorporated an updated generic classification for Heuchereae based on previous results (Folk et al. 2021). Greatly strengthened taxonomic sampling has confirmed previous results and provided additional support for the segregation of *Mitella* into multiple genera (*Mitella, Mitellastra, Pectiantia, Ozomelis, Spuriomitella, Asimitellaria, Brewerimitella*; Folk et al. 2021). While some relationships, such as the placement of *Heuchera* relative to the *Ozomelis* group, remain uncertain (Fig. 2), the backbone of Heuchereae has largely been resolved. Relationships within genera typically receive decisive support and are consistent across analyses. Important exceptions are within *Heuchera*: *H. bracteata, H. hallii,* and *H. caespitosa* had unconventional placements in the concatenation analysis. Coalescent and concatenated analyses also differed in the position of *H. glabra,* whether as sister to all other species of *Heuchera* (concatenation, with weak support) or in a position close to *H. micrantha* (coalescence, with decisive support). Both of these placements have been seen previously in different locus sets on the same accessions (Folk and Freudenstein 2014; Folk et al. 2017). The inconsistency of placement for these taxa may represent two alternative placements in the gene tree set, resolved differently depending on analytical method and dataset, with the placement in coalescent analyses most consistent with previous work. The position of *H. glabra* and resolution within *Heuchera* sect. *Rhodoheuchera* have remained difficult despite repeated phylogenomic study, and a more focused study of these taxa is a prime target for future work in the group.

### Hybrid detection and distribution

With more taxon sampling, we found support for a previous hypothesis of hybridization that was tentative given a lack of outgroup sampling (Folk et al. 2017), as well as other hypotheses in the literature (Soltis et al. 1991; Okuyama et al. 2005). This work identified a new chloroplast capture event between *Lithophragma* and *Tolmiea* (Fig. 4), genera of western North America that do not resemble each other morphologically and are not sister. While both genera contain polyploid species or populations, the entire *Tolmiea* clade is embedded within *Lithophragma*, suggesting the hybridization occurred in a diploid ancestor. This accords with the chromosome survey and phylogeny of Folk and Freudenstein (2014) focused on *Heuchera*, which showed that polyploidy was restricted to individual species or (more commonly) populations of species. There are no instances of fixed polyploidy in clades treated above the species level for the Heuchereae of North America, and most known cases of polyploidy do not align with chloroplast capture events, suggesting autopolyploidy as the primary mechanism of polyploidization in Heuchereae and an unclear relationship (if any) to hybridization. The exceptions are three examples of allopolyploidy known so far: *Heuchera rubescens* var. *cuneata* (Folk et al. 2017), some populations of *Lithophragma bolanderi* (Kuzoff et al. 1999), and the Asian clade *Asimitellaria* (Okuyama et al. 2012).

Our focus has been on a specific hybridization process: chloroplast capture as evident from the comparison of nuclear and chloroplast DNA sequence data. Chloroplast capture may be the most commonly reported form of reported hybridization in plants in terms of the number of evolutionary events detected so far, although it remains uncertain whether this only reflects its more straightforward detection (Sambatti et al., 2008, p. 1089). In Heuchereae, deep intergeneric conflict is less evident when comparing among only nuclear gene trees (see Discussion in Folk et al. 2017) and does not reflect the phylogenetic signal recovered from chloroplast data. This justifies our use of a single species tree estimate rather than a network for ancestral reconstruction. The small weight of extra hybridization edges in the context of introgression (see Bastide et al. 2018) or the involvement of chloroplast housekeeping loci that are unlikely to underlie niche traits (Hahn and Nakhleh 2016) could limit the impact of hybridization on ancestral reconstruction. While feeling justified in constraining our hybridization focus to the chloroplast genome, we have not completely leveraged the potential of our data to reveal further potential hybrid that could be evident from nuclear data.

## Conclusions

We have found evidence supporting a classic hypothesis that the mid- to late Pleistocene paleoclimates were potential drivers of hybridization, the results of which are observed in some present-day floras. Glacial cool periods represent opportunities for hybridization by causing large-scale range shifts in plant communities and contraction in the distributions of individual species, which would have created novel patterns of sympatry and parapatry among plant lineages that ancestrally lacked opportunities for gene flow. We found evidence that most chloroplast capture events in Heuchereae date to the mid- to late Pleistocene, concomitant with clade-wide range restriction during Pleistocene cool conditions, indicating a role for past climate change in promoting prolific chloroplast capture in the clade. The ancestral reconstruction approach we implement here will have broad applicability for testing questions about niche biology in deep time.

## Acknowledgements

This work was supported by DBI-1523667 (to R.A.F.), DBI-1458640 and DBI-1930007 (to D.E.S. and P.S.S.), iDigBio (DBI-1547229 to P.S.S.), and DGE-1842473 (to M.L.G.). C. Siniscalchi is thanked for input on a draft version of this manuscript. We thank J.N. Thompson for samples and feedback on this manuscript.

## Notes

### Competing Interest Statement

The authors have declared no competing interest.

