## Supplement for "Identifying climatic drivers of hybridization in Heuchereae (Saxifragaceae)"

### Supplemental Figures

**Fig. S1.** Topology recovered in RAxML using near-complete chloroplast genomes. Branch labels represent bootstrap percentages ≥ 50%. Branch lengths represent per-site substitutions. Informal chloroplast clade names from Folk et al. (2017) are labeled with gray boxes. Outgroup branches (*Darmera, Rodgersia, Peltoboykinia*) are omitted for compact representation.

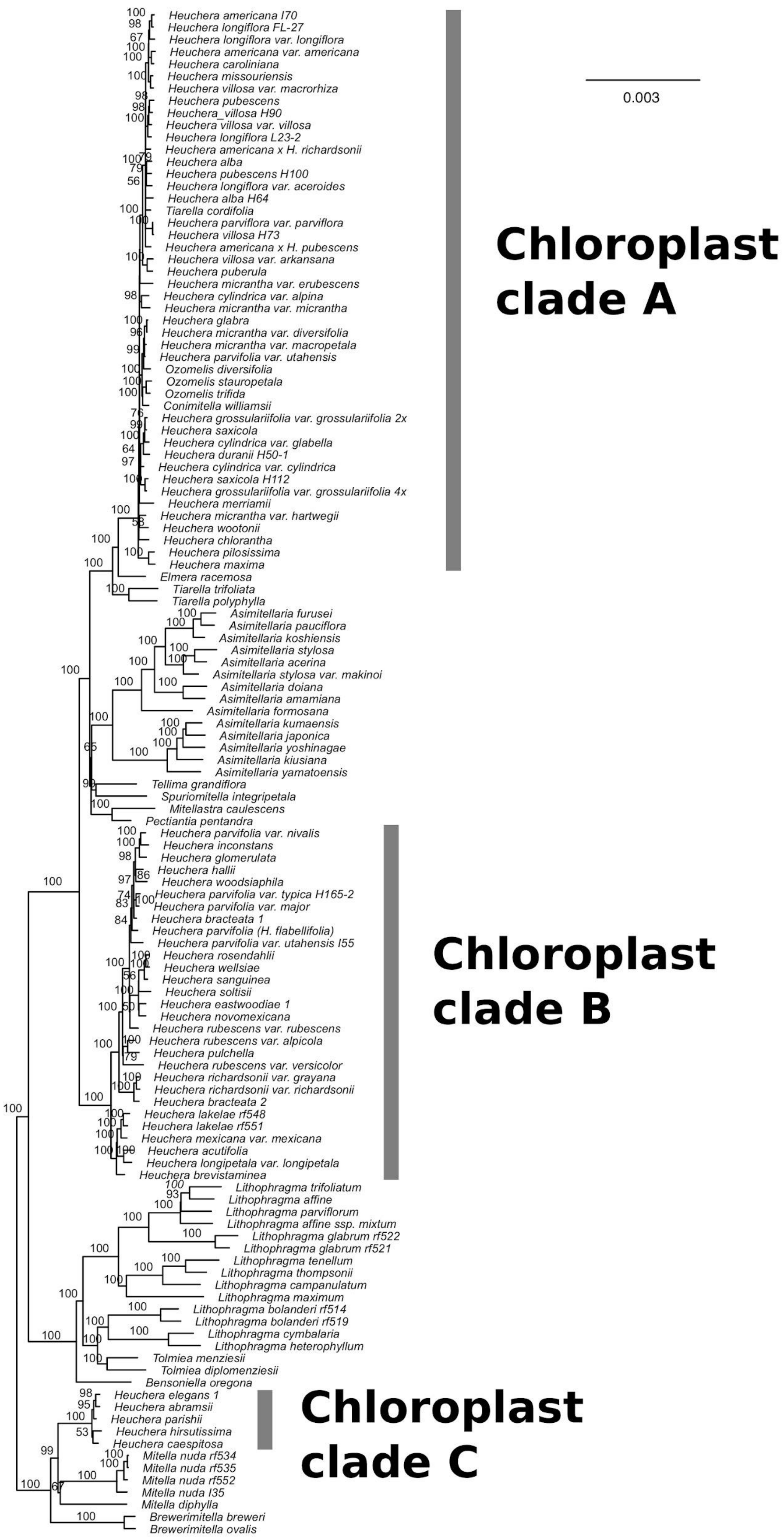
0.003
Chloroplast clade A
Chloroplast clade B
Chloroplast clade C
Heuchera americana I70
Heuchera longiflora FL-27
Heuchera longiflora var. longiflora
Heuchera americana var. americana
Heuchera caroliniana
Heuchera missouriensis
Heuchera villosa var. macrorhiza
Heuchera pubescens
Heuchera_villosa H90
Heuchera villosa var. villosa
Heuchera longiflora L23-2
Heuchera americana x H. richardsonii
Heuchera alba
Heuchera pubescens H100
Heuchera longiflora var. aceroides
Heuchera alba H64
Tiarella cordifolia
Heuchera parviflora var. parviflora
Heuchera villosa H73
Heuchera americana x H. pubescens
Heuchera villosa var. arkansana
Heuchera puberula
Heuchera micrantha var. erubescens
Heuchera cylindrica var. alpina
Heuchera micrantha var. micrantha
Heuchera glabra
Heuchera micrantha var. diversifolia
Heuchera micrantha var. macropetala
Heuchera parvifolia var. utahensis
Ozomelis diversifolia
Ozomelis stauropetala
Ozomelis trifida
Conimitella williamsii
Heuchera grossulariifolia var. grossulariifolia 2x
Heuchera saxicola
Heuchera cylindrica var. glabella
Heuchera duranii H50-1
Heuchera cylindrica var. cylindrica
Heuchera saxicola H112
Heuchera grossulariifolia var. grossulariifolia 4x
Heuchera merriamii
Heuchera micrantha var. hartwegii
Heuchera wootonii
Heuchera chlorantha
Heuchera pilosissima
Heuchera maxima
Elmera racemosa
Tiarella trifoliata
Tiarella polyphylla
Asimitellaria furusei
Asimitellaria pauciflora
Asimitellaria koshiensis
Asimitellaria stylosa
Asimitellaria acerina
Asimitellaria stylosa var. makinoi
Asimitellaria doiana
Asimitellaria amamiana
Asimitellaria formosana
Asimitellaria kumaensis
Asimitellaria japonica
Asimitellaria yoshinagae
Asimitellaria kiusiana
Asimitellaria yamatoensis
Tellima grandiflora
Spuriomitella integripetala
Mitellastra caulescens
Pectiantia pentandra
Heuchera parvifolia var. nivalis
Heuchera inconstans
Heuchera glomerulata
Heuchera hallii
Heuchera woodsiaphila
Heuchera parvifolia var. typica H165-2
Heuchera parvifolia var. major
Heuchera bracteata 1
Heuchera parvifolia (H. flabellifolia)
Heuchera parvifolia var. utahensis I55
Heuchera rosendahlii
Heuchera wellsiae
Heuchera sanguinea
Heuchera soltisii
Heuchera eastwoodiae 1
Heuchera novomexicana
Heuchera rubescens var. rubescens
Heuchera rubescens var. alpicola
Heuchera pulchella
Heuchera rubescens var. versicolor
Heuchera richardsonii var. grayana
Heuchera richardsonii var. richardsonii
Heuchera bracteata 2
Heuchera lakelae rf548
Heuchera lakelae rf551
Heuchera mexicana var. mexicana
Heuchera acutifolia
Heuchera longipetala var. longipetala
Heuchera brevistaminea
Lithophragma trifoliatum
Lithophragma affine
Lithophragma parviflorum
Lithophragma affine ssp. mixtum
Lithophragma glabrum rf522
Lithophragma glabrum rf521
Lithophragma tenellum
Lithophragma thompsonii
Lithophragma campanulatum
Lithophragma maximum
Lithophragma bolanderi rf514
Lithophragma bolanderi rf519
Lithophragma cymbalaria
Lithophragma heterophyllum
Tolmiea menziesii
Tolmiea diplomenziesii
Bensoniella oregona
Heuchera elegans 1
Heuchera abramsii
Heuchera parishii
Heuchera hirsutissima
Heuchera caespitosa
Mitella nuda rf534
Mitella nuda rf535
Mitella nuda rf552
Mitella nuda I35
Mitella diphylla
Brewerimitella breweri
Brewerimitella ovalis

**Fig. S2.** Dating result for the nuclear phylogeny from MCMCTREE. The x-axis represents time in millions of years; node bars represent 95% credibility intervals. Outgroup taxa are included here (cf. Figs. 2, 3).

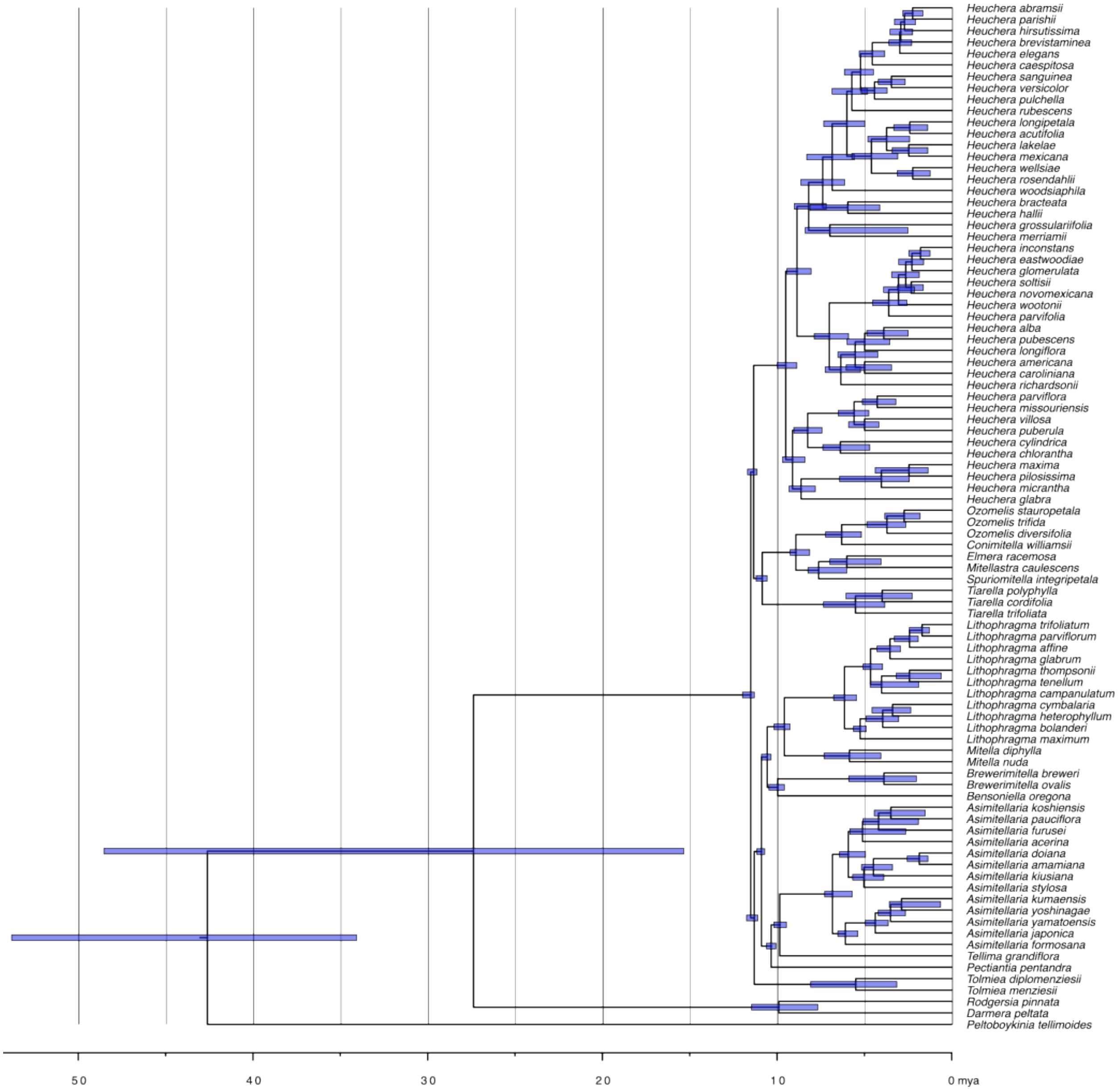

**Fig. S3.** Ancestral reconstructions across the 12 predictor variables on the ASTRAL topology. Given difficulties visualizing histogram data on trees, branches are colored by the bin with maximum probability density for each variable. Branch colors are on a spectrum from blue (low values) to red (high values), scaled to the minimum and maximum per variable. The original reconstructions and further plot data are at https://github.com/ryanafolk/heuchera_ancestral_niche/.

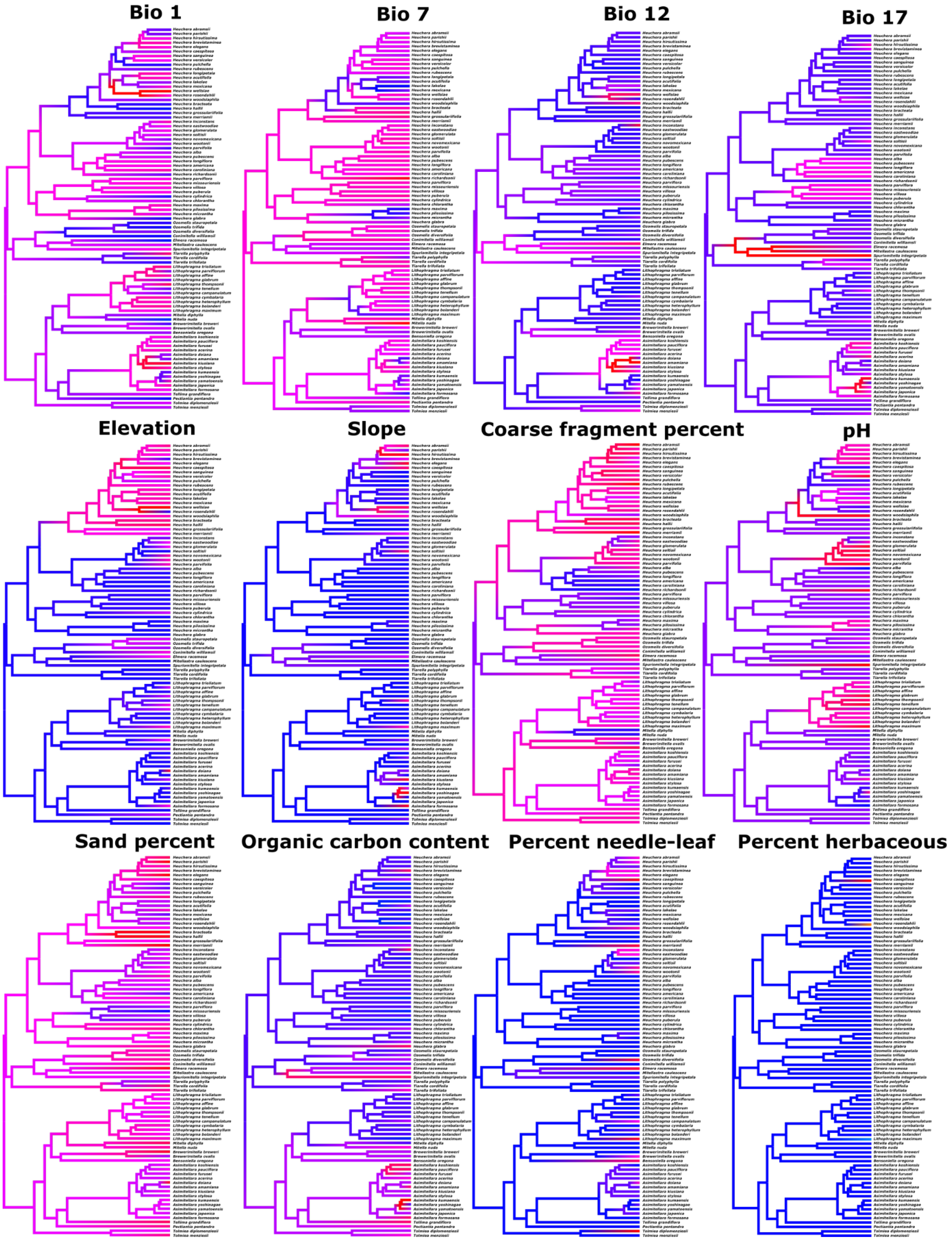

Bio 1
Bio 7
Bio 12
Bio 17
Elevation
Slope
Coarse fragment percent
pH
Sand percent
Organic carbon content
Percent needle-leaf
Percent herbaceous

**Fig. S4.** Chloroplast clade frequencies expected given the nuclear gene tree distribution and ILS alone. These simulations used a nuclear scaling factor of 2. The scale represents chloroplast genome per-site substitutions.

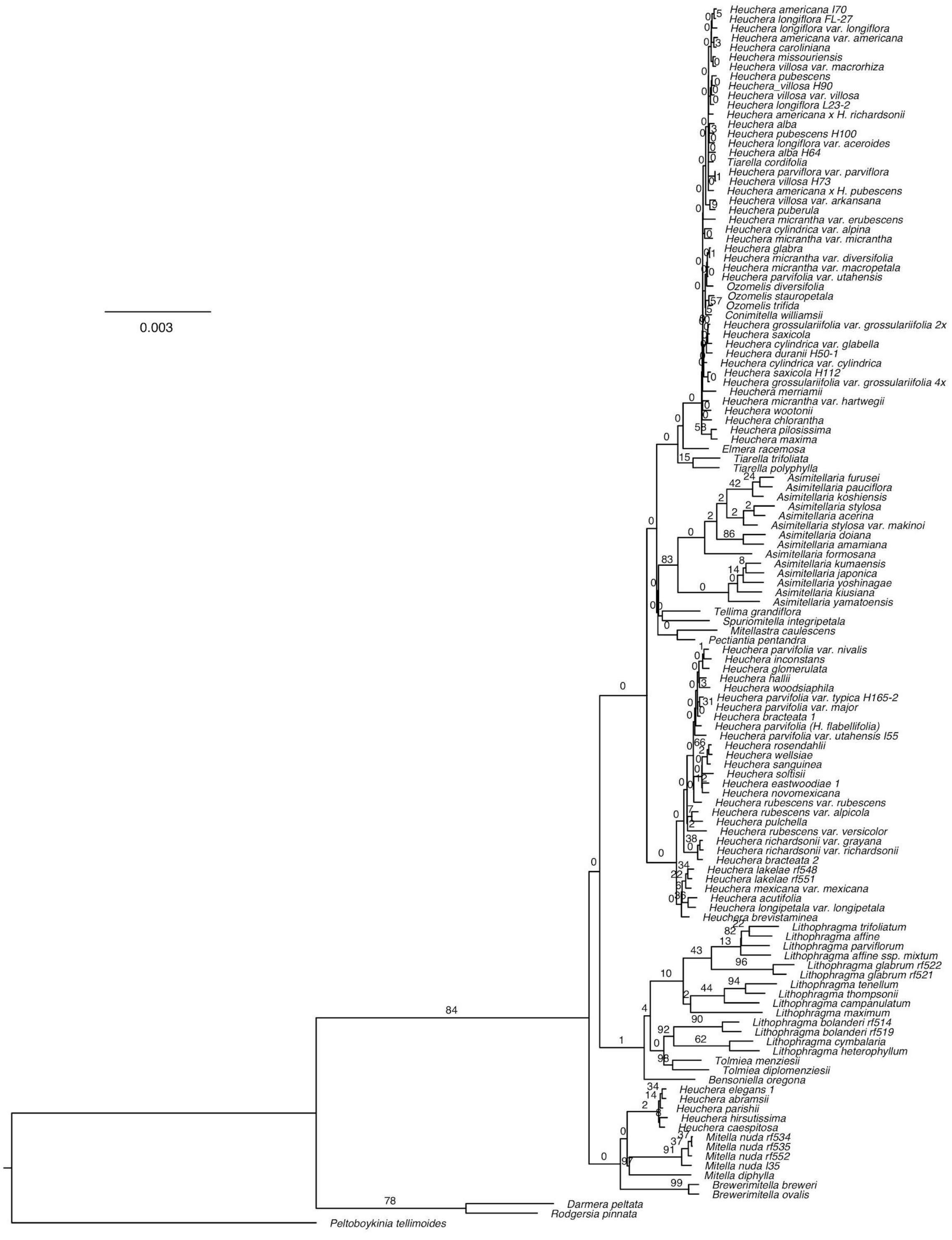
0.003
Heuchera americana I70
Heuchera longiflora FL-27
Heuchera longiflora var. longiflora
Heuchera americana var. americana
Heuchera caroliniana
Heuchera missouriensis
Heuchera villosa var. macrorhiza
Heuchera pubescens
Heuchera_villosa H90
Heuchera villosa var. villosa
Heuchera longiflora L23-2
Heuchera americana x H. richardsonii
Heuchera alba
Heuchera pubescens H100
Heuchera longiflora var. aceroides
Heuchera alba H64
Tiarella cordifolia
Heuchera parviflora var. parviflora
Heuchera villosa H73
Heuchera americana x H. pubescens
Heuchera villosa var. arkansana
Heuchera puberula
Heuchera micrantha var. erubescens
Heuchera cylindrica var. alpina
Heuchera micrantha var. micrantha
Heuchera glabra
Heuchera micrantha var. diversifolia
Heuchera micrantha var. macropetala
Heuchera parvifolia var. utahensis
Ozomelis diversifolia
Ozomelis stauropetala
Ozomelis trifida
Conimitella williamsii
Heuchera grossulariifolia var. grossulariifolia 2x
Heuchera saxicola
Heuchera cylindrica var. glabella
Heuchera duranii H50-1
Heuchera cylindrica var. cylindrica
Heuchera saxicola H112
Heuchera grossulariifolia var. grossulariifolia 4x
Heuchera merriamii
Heuchera micrantha var. hartwegii
Heuchera wootonii
Heuchera chlorantha
Heuchera pilosissima
Heuchera maxima
Elmera racemosa
Tiarella trifoliata
Tiarella polyphylla
Asimitellaria furusei
Asimitellaria pauciflora
Asimitellaria koshiensis
Asimitellaria stylosa
Asimitellaria acerina
Asimitellaria stylosa var. makinoi
Asimitellaria doiana
Asimitellaria amamiana
Asimitellaria formosana
Asimitellaria kumaensis
Asimitellaria japonica
Asimitellaria yoshinagae
Asimitellaria kiusiana
Asimitellaria yamatoensis
Tellima grandiflora
Spuriomitella integripetala
Mitellastra caulescens
Pectiantia pentandra
Heuchera parvifolia var. nivalis
Heuchera inconstans
Heuchera glomerulata
Heuchera hallii
Heuchera woodsiaphila
Heuchera parvifolia var. typica H165-2
Heuchera parvifolia var. major
Heuchera bracteata 1
Heuchera parvifolia (H. flabellifolia)
Heuchera parvifolia var. utahensis I55
Heuchera rosendahlii
Heuchera wellsiae
Heuchera sanguinea
Heuchera soltisii
Heuchera eastwoodiae 1
Heuchera novomexicana
Heuchera rubescens var. rubescens
Heuchera rubescens var. alpicola
Heuchera pulchella
Heuchera rubescens var. versicolor
Heuchera richardsonii var. grayana
Heuchera richardsonii var. richardsonii
Heuchera bracteata 2
Heuchera lakelae rf548
Heuchera lakelae rf551
Heuchera mexicana var. mexicana
Heuchera acutifolia
Heuchera longipetala var. longipetala
Heuchera brevistaminea
Lithophragma trifoliatum
Lithophragma affine
Lithophragma parviflorum
Lithophragma affine ssp. mixtum
Lithophragma glabrum rf522
Lithophragma glabrum rf521
Lithophragma tenellum
Lithophragma thompsonii
Lithophragma campanulatum
Lithophragma maximum
Lithophragma bolanderi rf514
Lithophragma bolanderi rf519
Lithophragma cymbalaria
Lithophragma heterophyllum
Tolmiea menziesii
Tolmiea diplomenziesii
Bensoniella oregona
Heuchera elegans 1
Heuchera abramsii
Heuchera parishii
Heuchera hirsutissima
Heuchera caespitosa
Mitella nuda rf534
Mitella nuda rf535
Mitella nuda rf552
Mitella nuda I35
Mitella diphylla
Brewerimitella breweri
Brewerimitella ovalis
Darmera peltata
Rodgersia pinnata
Peltoboykinia tellimoides

**Fig. S5.** Chloroplast clade frequencies expected given the nuclear gene tree distribution and ILS alone. These simulations used a nuclear scaling factor of 4. The scale represents chloroplast genome per-site substitutions.

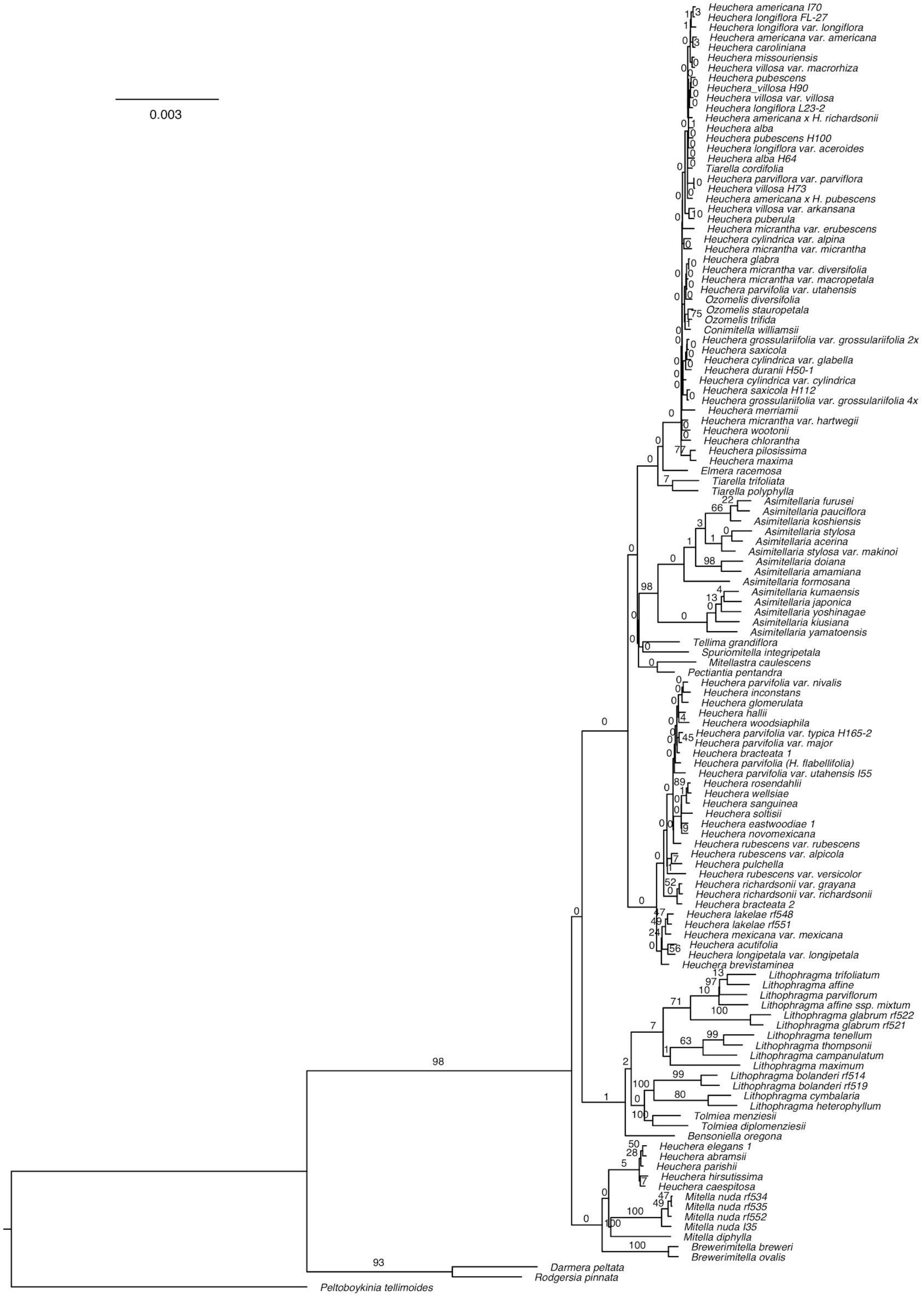
0.003
Heuchera americana I70
Heuchera longiflora FL-27
Heuchera longiflora var. longiflora
Heuchera americana var. americana
Heuchera caroliniana
Heuchera missouriensis
Heuchera villosa var. macrorhiza
Heuchera pubescens
Heuchera_villosa H90
Heuchera villosa var. villosa
Heuchera longiflora L23-2
Heuchera americana x H. richardsonii
Heuchera alba
Heuchera pubescens H100
Heuchera longiflora var. aceroides
Heuchera alba H64
Tiarella cordifolia
Heuchera parviflora var. parviflora
Heuchera villosa H73
Heuchera americana x H. pubescens
Heuchera villosa var. arkansana
Heuchera puberula
Heuchera micrantha var. erubescens
Heuchera cylindrica var. alpina
Heuchera micrantha var. micrantha
Heuchera glabra
Heuchera micrantha var. diversifolia
Heuchera micrantha var. macropetala
Heuchera parvifolia var. utahensis
Ozomelis diversifolia
Ozomelis stauropetala
Ozomelis trifida
Conimitella williamsii
Heuchera grossulariifolia var. grossulariifolia 2x
Heuchera saxicola
Heuchera cylindrica var. glabella
Heuchera duranii H50-1
Heuchera cylindrica var. cylindrica
Heuchera saxicola H112
Heuchera grossulariifolia var. grossulariifolia 4x
Heuchera merriamii
Heuchera micrantha var. hartwegii
Heuchera wootonii
Heuchera chlorantha
Heuchera pilosissima
Heuchera maxima
Elmera racemosa
Tiarella trifoliata
Tiarella polyphylla
Asimitellaria furusei
Asimitellaria pauciflora
Asimitellaria koshiensis
Asimitellaria stylosa
Asimitellaria acerina
Asimitellaria stylosa var. makinoi
Asimitellaria doiana
Asimitellaria amamiana
Asimitellaria formosana
Asimitellaria kumaensis
Asimitellaria japonica
Asimitellaria yoshinagae
Asimitellaria kiusiana
Asimitellaria yamatoensis
Tellima grandiflora
Spuriomitella integripetala
Mitellastra caulescens
Pectiantia pentandra
Heuchera parvifolia var. nivalis
Heuchera inconstans
Heuchera glomerulata
Heuchera hallii
Heuchera woodsiaphila
Heuchera parvifolia var. typica H165-2
Heuchera parvifolia var. major
Heuchera bracteata 1
Heuchera parvifolia (H. flabellifolia)
Heuchera parvifolia var. utahensis I55
Heuchera rosendahlii
Heuchera wellsiae
Heuchera sanguinea
Heuchera soltisii
Heuchera eastwoodiae 1
Heuchera novomexicana
Heuchera rubescens var. rubescens
Heuchera rubescens var. alpicola
Heuchera pulchella
Heuchera rubescens var. versicolor
Heuchera richardsonii var. grayana
Heuchera richardsonii var. richardsonii
Heuchera bracteata 2
Heuchera lakelae rf548
Heuchera lakelae rf551
Heuchera mexicana var. mexicana
Heuchera acutifolia
Heuchera longipetala var. longipetala
Heuchera brevistaminea
Lithophragma trifoliatum
Lithophragma affine
Lithophragma parviflorum
Lithophragma affine ssp. mixtum
Lithophragma glabrum rf522
Lithophragma glabrum rf521
Lithophragma tenellum
Lithophragma thompsonii
Lithophragma campanulatum
Lithophragma maximum
Lithophragma bolanderi rf514
Lithophragma bolanderi rf519
Lithophragma cymbalaria
Lithophragma heterophyllum
Tolmiea menziesii
Tolmiea diplomenziesii
Bensoniella oregona
Heuchera elegans 1
Heuchera abramsii
Heuchera parishii
Heuchera hirsutissima
Heuchera caespitosa
Mitella nuda rf534
Mitella nuda rf535
Mitella nuda rf552
Mitella nuda I35
Mitella diphylla
Brewerimitella breweri
Brewerimitella ovalis
Darmera peltata
Rodgersia pinnata
Peltoboykinia tellimoides

### Supplemental Tables

**Table S1.** Accessions included in this study.

| Accession | Species | Source | Collector | Collector number | Voucher herbarium location | Locality |
|---|---|---|---|---|---|---|
| I23 | *Asimitellaria acerina* | Newly sequenced | Okuyama | MA-MY1 | Field collection | Yohoro, Maizuru City, Kyoto Pref., Japan |
| I114 | *Asimitellaria amamiana* | Newly sequenced | Okuyama | MAM-02 | Field collection | Sumiyo-cho, Amami city, Kagoshima pref., Japan |
| I116 | *Asimitellaria doiana* | Newly sequenced | Umemoto | MD1501 | Living accession, Tsukuba Botanical Garden | In the vicinity of Miyanoura river, Yakushima Isl. Kagoshima pref. Japan |
| I117 | *Asimitellaria formosana* | Newly sequenced | Okuyama | MFO-LL31 | Living accession, Tsukuba Botanical Garden | Lala-shan, Taoyuan Hsien, Taiwan |
| I39 | *Asimitellaria furusei var. subramosa* | Folk et al. 2018a | Okuyama | MSU-Ko4 | Living accession, Tsukuba Botanical Garden | Jorakuji, Koryo-cho, Shimane Pref., Japan |
| I37 | *Asimitellaria japonica* | Newly sequenced | Okuyama | MJ-T5 | Living accession, Tsukuba Botanical Garden | Takachiho-kyo, Takachiho-cho, Miyazaki Pref., Japan |
| I120 | *Asimitellaria kiusiana* | Newly sequenced | Okuyama | MK-I16 | Living accession, Tsukuba Botanical Garden | Oh-taki, Itsuki-mura, Kumamoto pref., Japan |
| I38 | *Asimitellaria koshiensis* | Newly sequenced | Okuyama | MKO-M0901 | Living accession, Tsukuba Botanical Garden | Igashima, Aga-cho, Niigata Pref., JAPAN |
| I121 | *Asimitellaria kumaensis* | Newly sequenced | Okuyama | MY-I1302 | Living accession, | Oh-taki, Itsuki-mura, Kumamoto pref., Japan |

| | | | | | | |
|---|---|---|---|---|---|---|
| | | | | | Tsukuba Botanical Garden | |
| I36 | *Asimitellaria pauciflora* | Folk et al. 2018a | Okuyama | MP−K1004 | Living accession, Tsukuba Botanical Garden | Kibune-okumiya, Sakyo, Kyoto City, Kyoto Pref., Japan |
| I31 | *Asimitellaria stylosa* var. *makinoi* | Newly sequenced | Okuyama | MSM-KKW2 | Living accession, Tsukuba Botanical Garden | Kaikawa, Naka-cho, Tokushima Pref, Japan |
| I26 | *Asimitellaria stylosa* var. *stylosa* | Folk et al. 2018a | Okuyama | MSS-KZ1106 | Living accession, Tsukuba Botanical Garden | Kosaka, Ibigawa-cho, Gifu Pref., Japan |
| I119 | *Asimitellaria yamatoensis* | Newly sequenced | Okuyama | MJ-A1107 | Living accession, Tsukuba Botanical Garden | Akame 48 falls, Nabari city, Mie pref., Japan |
| I22 | *Asimitellaria yoshinagae* | Newly sequenced | Sakaguchi | MY-KW1 | Living accession, Tsukuba Botanical Garden | Naka-cho, Tokushima Pref., Japan |
| H148 | *Bensoniella oregona* | Folk et al. 2017 | Folk | 148 | OS | FS Rd 23, near Bear Camp, Siskiyou Mts, OR, USA |
| I33 | *Brewerimitella breweri* | Folk et al. 2018a | Okuyama | MB-RL9 | Field collection | Rainy Lake Pass, Chelan Co., Washington, USA |
| I32 | *Brewerimitella ovalis* | Folk et al. 2018a | Okuyama | MO-FC8 | Living accession, Tsukuba Botanical Garden | Fletcher Canyon, Jefferson Co., Washington, USA |
| H109 | *Conimitella williamsii* | Newly sequenced | Folk | 109 | OS | Hellroaring Trail, Spanish Creek Basin, MT, USA |
| I48 | *Darmera peltata* | Newly sequenced | Folk | I-48 | OS | Cultivated |
| I89 | *Elmera racemosa* | Folk et al. 2018a | Soltis and Soltis | 2179 | WS | Kittitas Co., Washington, USA |
| H45 | *Heuchera abramsii* | Folk et al. 2017 | Folk | 45 | OS | Mount Baldy, CA, USA |

| | | | | | | |
|---|---|---|---|---|---|---|
| I74 | *Heuchera acutifolia* | Folk et al. 2017 | Matus | 14560 | TEX | Rancho Poza, VE, Mexico |
| H63 | *Heuchera alba* | Folk et al. 2017 | Folk | 63 | OS | Along Hwy 55 3 mi W of Onego, WV, USA |
| H64 | *Heuchera alba* | Folk et al. 2018b | Folk | 64 | OS | Pendleton County, WV, USA |
| I70 | *Heuchera americana* | Folk et al. 2018b | Folk | I-70 | OS | Polk County, TN, USA |
| H70 | *Heuchera americana var. americana* | Folk et al. 2017 | Folk | 70 | OS | Berry Road, Wayne National Forest, OH, USA |
| H104 | *Heuchera americana × H. pubescens* | Folk et al. 2017 | Folk | 104 | OS | Bank near Sandstone Falls, WV, USA |
| H59 | *Heuchera americana × H. richardsonii* | Folk et al. 2017 | Folk | 59 | OS | Roadside woods off Hwy 19, Mark Twain NF, Shannon Co. MO, USA |
| H52 | *Heuchera bracteata* | Folk et al. 2017 | Folk | 52 | OS | Dowdy Lake, CO, USA |
| H164 | *Heuchera bracteata* | Newly sequenced | Folk | 164 | OS | Blair-Wallis road near campground, Medicine Bow National Forest, WY, USA |
| H37 | *Heuchera brevistaminea* | Folk et al. 2017 | Folk | 37 | OS | Side trail near Pacific Crest Trail, Laguna Mts., CA, USA |
| H48 | *Heuchera caespitosa* | Folk et al. 2017 | Folk | 48 | OS | Rose Lake Falls, CA, USA |
| H216 | *Heuchera caroliniana* | Folk et al. 2017 | Folk | 216 | OS | Bank along Rocky Creek, intersection with Hwy 97, SC, USA |
| H138 | *Heuchera chlorantha* | Folk et al. 2017 | Folk | 138 | OS | Along Hwy 35, OR, USA |
| H108 | *Heuchera cylindrica* ("*H. saxicola*") | Folk et al. 2017 | Folk | 108 | OS | Hellroaring Trail, Spanish Creek Basin, MT, USA |
| H112 | *Heuchera cylindrica* ("*H. saxicola*") | Newly sequenced | Folk | 112 | OS | Galena Gulch road, just off Hwy 15, MT, USA |
| H140 | *Heuchera cylindrica var. alpina* | Folk et al. 2017 | Folk | 140 | OS | Cliffs above Deschutes River, OR, USA |
| H159 | *Heuchera cylindrica var. cylindrica* | Folk et al. 2017 | Folk | 159 | OS | Dry banks above Salmon River Road, ID, USA |
| I57 | *Heuchera cylindrica var. glabella* | Folk et al. 2017 | Folk | I-57 | OS | Fishtrap Lake, WA, USA |
| H35 | *Heuchera eastwoodiae* | Folk et al. 2017 | Folk | 35 | OS | Small canyon along Senator Hwy, 3.6 mi S of Groom Creek, AZ, USA |
| H44 | *Heuchera elegans* | Folk et al. 2017 | Folk | 44 | OS | Talus above road to Crystal Lake, 0.5 mi from Hwy 39, CA, USA |
| I46 | *Heuchera glabra* | Folk et al. 2017 | Folk | I-46 | OS | Cliffs along Glacier Highway near Juneau, AK, USA |
| I5 | *Heuchera glomerulata* | Folk et al. 2017 | Folk | I-5 | OS | Cliffs along Mule Creek Road, NM near AZ border, USA |
| H154 | *Heuchera grossulariifolia* var. *grossulariifolia* 2x | Folk et al. 2017 | Folk | 154 | OS | Cliffs above Shaw Mountain Road near Boise, ID, USA |

| | | | | | | |
|---|---|---|---|---|---|---|
| H160 | *Heuchera grossulariifolia* var. *grossulariifolia 4x* | Folk et al. 2017 | Folk | 160 | OS | Cliff above roadside parking lot outside of Riggins, ID, USA |
| H56 | *Heuchera hallii* | Folk et al. 2017 | Folk | 56 | OS | Along Rock Creek Hills Road, Lost Park, CO, USA |
| H39 | *Heuchera hirsutissima* | Folk et al. 2017 | Folk | 39 | OS | Mount San Jacinto, CA, USA |
| H33 | *Heuchera inconstans* | Folk et al. 2017 | Folk | 33 | OS | West Fork Creek Canyon, AZ, USA |
| I126 | *Heuchera lakelae* | Newly sequenced | Hinton | 18331 | TEX | Sierra La Marta, Mpio. Arteaga, Coahuila, Mexico |
| I129 | *Heuchera lakelae* | Newly sequenced | Villarreal | 5471 | TEX | El Coahuilón, Sierra de la Marta, Mpio. Arteaga, Coahuila, Mexico |
| L23-2 | *Heuchera longiflora* | Folk et al. 2018b | Folk | L-23 | OS | Pike County, KY, USA |
| FL-27 | *Heuchera longiflora* | Folk et al. 2018b | Floden | s.n. | TENN | Talledega County, AL, USA |
| H65 | *Heuchera longiflora* var. *longiflora* | Folk et al. 2017 | Folk | 65 | OS | Natural Bridge State Park, KY, USA |
| H98 | *Heuchera longiflora* var. *longiflora* | Folk et al. 2017 | Folk | 98 | OS | Rocky bank, Hwy 209 ~5 mi N of Paint Rock, NC, USA |
| I21 | *Heuchera longipetala* var. *longipetala* | Folk et al. 2017 | Folk | I-21 | OS | Sierra de Tepoztlan, MO, Mexico |
| I6 | *Heuchera maxima* | Folk et al. 2017 | Folk | I-6 | OS | 0.5 mi SE of Stanton Ranch, Santa Cruz Island, CA, USA |
| H150 | *Heuchera merriamii* | Folk et al. 2017 | Folk | 150 | OS | Outcrops above Little Castle Lake, CA, USA |
| I51 | *Heuchera mexicana* var. *mexicana* | Folk et al. 2017 | Folk | I-51 | OS | Bufa del Diente, Sierra Chiquita, Tamaulipas, NL, Mexico |
| H134 | *Heuchera micrantha* var. *diversifolia* | Folk et al. 2017 | Folk | 134 | OS | Dry cliffs along Hwy 542 near Excelsior Camp, WA, USA |
| H49 | *Heuchera micrantha* var. *erubescens* | Folk et al. 2017 | Folk | 49 | OS | Roadside outcrops on Hwy 41, Inyo Co., CA, USA |
| H145 | *Heuchera micrantha* var. *hartwegii* | Folk et al. 2017 | Folk | 145 | OS | Cliff along Hwy 101 (side of Humbug Mountain), OR, USA |
| H146 | *Heuchera micrantha* var. *macropetala* | Folk et al. 2017 | Folk | 146 | OS | FS 2308 1 mi from Rd 23, Siskiyou Mts., OR, USA |
| H141 | *Heuchera micrantha* var. *micrantha* | Folk et al. 2017 | Folk | 141 | OS | Viento State Park, OR, USA |
| I69 | *Heuchera missouriensis* | Folk et al. 2017 | Folk | I-69 | OS | Little Grand Canyon near Carbondale, IL, USA |
| H24 | *Heuchera novomexicana* | Folk et al. 2017 | Folk | 24 | OS | Emory Pass Trail, NM, USA |
| H43 | *Heuchera parishii* | Folk et al. 2017 | Folk | 43 | OS | Sugarloaf Mountain, CA, USA |
| H72 | *Heuchera parviflora* var. *parviflora* | Folk et al. 2017 | Fok | 72 | OS | Natural Bridge State Park, KY, USA |

| | | | | | | |
|---|---|---|---|---|---|---|
| H50-1 | *Heuchera parvifolia* ("*H. duranii*") | Newly sequenced | Folk | 50 | OS | McAfee Meadow, White Mountains, Mono County, CA, USA |
| H120 | *Heuchera parvifolia* ("*H. flabellifolia*") | Folk et al. 2017 | Folk | 120 | OS | Dry talus slope, Glacier Co., MT, USA |
| H53 | *Heuchera parvifolia* var. *major* | Folk et al. 2017 | Folk | 53 | OS | Poudre Canyon Hwy, CO, USA |
| H55 | *Heuchera parvifolia* var. *nivalis* | Folk et al. 2017 | Folk | 55 | OS | Along Rock Creek Hills Road, Lost Park, CO, USA |
| H165-2 | *Heuchera parvifolia* var. *typica* | Newly sequenced | Folk | 165 | OS | Vedauwoo Glen road near Hwy 80, vicinity of campground, Medicine Bow National Forest, WY, USA |
| I56 | *Heuchera parvifolia* var. *utahensis* | Folk et al. 2017 | Folk | I-56 | OS | Cotterel Mountains., ID, USA |
| I55 | *Heuchera parvifolia* var. *utahensis* | Newly sequenced | Folk | I-55 | OS | Rock Butte NA, Wasatch National Forest, UT, USA |
| I59 | *Heuchera pilosissima* | Folk et al. 2017 | Folk | I-59 | OS | Marshall, Marin Co., CA, USA |
| H69 | *Heuchera puberula* | Folk et al. 2017 | Folk | 69 | OS | Pea Vine Road near Big Springs, Ozark National Riverways, MO, USA |
| H96 | *Heuchera pubescens* | Folk et al. 2017 | Folk | 96 | OS | Pilot Mountain State Park, NC, USA |
| H100 | *Heuchera pubescens* | Folk et al. 2018b | Folk | 100 | OS | Floyd County, VA, USA |
| H20 | *Heuchera pulchella* | Folk et al. 2017 | Folk | 20 | OS | Sandia Crest, NM, USA |
| H166 | *Heuchera richardsonii* var. *grayana* | Folk et al. 2017 | Folk | 166 | OS | Roadside near Ledges State Park, IA, USA |
| I58 | *Heuchera richardsonii* var. *richardsonii* | Folk et al. 2017 | Folk | I-58 | OS | Wainwright, Alberta, Canada |
| I9 | *Heuchera rosendahlii* | Folk et al. 2017 | Gurney | s.n. | RSA | Basaseachic Falls, CH, Mexico |
| I43 | *Heuchera rubescens* var. *alpicola* | Folk et al. 2017 | Folk | I-43 | OS | Lone Pine Creek above Mirror Lake, Sierra Nevada Mts., CA, USA |
| I19 | *Heuchera rubescens* var. *rubescens* | Folk et al. 2017 | Folk | I-19 | OS | Monroe Canyon, E of Richfield, UT, USA |
| H26 | *Heuchera rubescens* var. *versicolor* | Folk et al. 2017 | Folk | 26 | OS | Outcrop above Hwy 159, NM, USA |
| I4 | *Heuchera sanguinea* | Folk et al. 2017 | Folk | I-4 | OS | Middlemarch Pass, Dragoon Mts., AZ, USA |
| I85 | *Heuchera soltisii* | Folk et al. 2017 | Folk | I-85 | OS | Aguirre Springs, Organ Mts., NM, USA |
| H192 | *Heuchera villosa* var. *arkansana* | Folk et al. 2017 | Folk | 192 | OS | Weddington Gap, AR, USA |
| H78 | *Heuchera villosa* var. *intermedia* | Folk et al. 2017 | Folk | 78 | OS | Long Lick Road, Shawnee State Forest, OH, USA |
| I122 | *Heuchera villosa* var. *macrorhiza* | Newly sequenced | Floden | s.n. | Field collection | Big Bottom Road along the Cumberland River north of Cookeville, TN, USA |

| | | | | | | |
|---|---|---|---|---|---|---|
| H90 | *Heuchera villosa* var. *villosa* | Newly sequenced | Folk | 90 | OS | 0.25 mi S of Lewis Fork Overlook on Blue Ridge Parkway, NC, USA |
| I1 | *Heuchera villosa* var. *villosa* | Folk et al. 2017 | Folk | I-1 | OS | Devil's Courthouse, NC, USA |
| I7 | *Heuchera wellsiae* | Folk et al. 2017 | Gentry | 7198 | RSA | Los Pucheros, Sierra Surotato, SI, Mexico |
| H23 | *Heuchera woodsiaphila* | Folk et al. 2017 | Folk | 23 | OS | Scree slope, Capitan Peak, NM, USA |
| H22 | *Heuchera wootonii* | Folk et al. 2017 | Folk | 22 | OS | Dry bank, Capitan Peak, NM, USA |
| rf523 | *Lithophragma affine* | Newly sequenced | Thompson | 9818 | Field collection | UC Hastings Reserve, CA |
| rf516 | *Lithophragma affine* ssp. *mixtum* | Newly sequenced | Thompson | 6904 | Field collection | Ranger Peak, CA |
| rf514 | *Lithophragma bolanderi* | Newly sequenced | Thompson | 4709 | Field collection | Marble Falls Trail, Sequoia National Park, CA |
| rf519 | *Lithophragma bolanderi* | Newly sequenced | Thompson | 8127 | Field collection | South Fork Merced River, CA |
| rf517 | *Lithophragma campanulatum* | Newly sequenced | Thompson | 7596 | Field collection | Pit River, CA |
| rf513 | *Lithophragma cymbalaria* | Newly sequenced | Thompson | 1741 | Field collection | Colson Canyon, CA |
| rf522 | *Lithophragma glabrum* | Newly sequenced | Thompson | 9292 | Field collection | Washington Hill, CA |
| rf521 | *Lithophragma glabrum* | Newly sequenced | Thompson | 8438 | Field collection | Klickitat River, WA |
| rf512 | *Lithophragma heterophyllum* | Newly sequenced | Thompson | 773 | Field collection | UC Hastings Reserve, CA |
| rf518 | *Lithophragma maximum* | Newly sequenced | Thompson | 7735-1 | Seed from Rancho Santa Ana Botanical Garden | San Clemente Island, CA |
| rf511 | *Lithophragma parviflorum* | Newly sequenced | Thompson | s.n. | Field collection | Keating Ridge, ID |
| rf524 | *Lithophragma tenellum* | Newly sequenced | Thompson | 10516 | Field collection | Buck Creek Overlook, CA |
| rf520 | *Lithophragma thompsonii* | Newly sequenced | Thompson | 8355 | Field collection | Ephrata, WA |
| rf515 | *Lithophragma trifoliatum* | Newly sequenced | Thompson | 6794 | Field collection | Hogsback Road, CA |
| H88 | *Mitella diphylla* | Folk et al. 2017 | Folk | 88 | OS | Piatt Park, Munroe Co., OH, USA |
| I123 | *Mitella nuda* | Newly sequenced | Garton | 1171 | NY | Bog east of Provincial Paper Mill, Port Arthur, Thunder Bay District, Ontario, Canada |

| | | | | | | |
|---|---|---|---|---|---|---|
| I124 | *Mitella nuda* | Newly sequenced | Boivin | 12536 | NY | A l'ouest du lac Iosegun, District de Jasper-Edson, Alberta, Canada |
| I125 | *Mitella nuda* | Newly sequenced | Freudenstein | 3040 | OS | Iosco County, Michigan, USA |
| I35 | *Mitella nuda* | Folk et al. 2018a | Okuyama | MN01 | Living accession, Tsukuba Botanical Garden | Murii Trail, Engaru City, Hokkaido Pref., Japan |
| I25 | *Mitellastra caulescens* | Folk et al. 2018a | Okuyama | MC-FDR3 | Field collection | Federation Forest State Park, King Co., Washington, USA |
| I130 | *Ozomelis diversifolia* | Newly sequenced | Soltis and Soltis | 2428 | WS | Josephine County, Oregon, USA |
| H161 | *Ozomelis stauropetala* | Folk et al. 2017 | Folk | 161 | OS | Roadside bank near Minnetonkia Cave, ID, USA |
| I118 | *Ozomelis trifida* | Newly sequenced | Okuyama | MT1601 | Living accession, Tsukuba Botanical Garden | Near Blewett Pass, Wenatchee Mountains, Chelan Co., WA, USA |
| H128 | *Pectiantia pentandra* | Folk et al. 2017 | Folk | 128 | OS | Pacific Crest Trail access from hwy 90, WA, USA |
| I49 | *Peltoboykinia tellimoides* | Newly sequenced | Folk | I-49 | OS | Cultivated |
| I50 | *Rodgersia pinnata* | Newly sequenced | Folk | I-50 | OS | Cultivated |
| I115 | *Spuriomitella integripetala* | Newly sequenced | Okuyama | MI-AZW1205 | Living accession, Tsukuba Botanical Garden | Hibara-wase-zawa, Kitashiobara-mura, Fukushima pref., Japan |
| H130:1 | *Tellima grandiflora* | Newly sequenced | Folk | 130 | OS | Glacier Creek road, Mount Baker, WA, USA |
| H89 | *Tiarella cordifolia* | Newly sequenced | Folk | 89 | OS | Hampton Hills Metropark, OH, USA |
| I113 | *Tiarella polyphylla* | Newly sequenced | Okuyama | s.n. | Living accession, Tsukuba Botanical Garden | Hibara-wase-zawa, Kitashiobara-mura, Fukushima pref., Japan |
| H127 | *Tiarella trifoliata* | Newly sequenced | Folk | 127 | OS | Pacific Crest Trail access from hwy 90, WA, USA |
| H131-C | *Tolmiea menziesii* | Newly sequenced | Folk | 131 | OS | Glacier Creek road, Mount Baker, WA, USA |
| H169:C | *Tomiea diplomenziesii* | Newly sequenced | Folk | 169 | OS | Small seep beside FS 23, Siskiyou National Forest, Josephine County, OR, USA |

**Table S2.** Comparison of dating results based on nuclear concatenation, nuclear coalescent, and chloroplast data. Shown are dates for named clades only, with – indicating clades were not recovered for the analysis in question; dates are crown dates; units are MYA (millions of years ago). Bracketed values represent 95% credibility intervals.

| **Clade** | **Concatenated nuclear tree** | **ASTRAL nuclear tree** | **Chloroplast tree** |
|---|---|---|---|
| *Asimitellaria* | 4.98 [3.56, 6.11] | 6.83 [5.70, 7.28] | 6.42 [5.60, 7.59] |
| *Brewerimitella* | 3.43 [0.7, 5.23] | 3.89 [2.03, 5.89] | 1.62 [0.9, 2.74] |
| *Heuchera* | 9.19 [7.86, 10.22] | 9.50 [8.87, 10.00] | – |
| *Heuchera* chloroplast clade A | – | – | 4.00 [3.44, 4.66] |
| *Heuchera* chloroplast clade B | – | – | 6.41 [5.63, 7.74] |
| *Heuchera* chloroplast clade C | – | – | 3.17 [1.59, 7.49] |
| Heuchereae | 11.99 [11.52, 13.12] | 11.51 [11.30, 11.98] | 10.20 [8.90, 13.23] |
| *Lithophragma* | 4.90 [4.09, 5.75] | 6.14 [5.45, 6.76] | – |
| *Mitella* | 5.64 [4.04, 7.02] | 5.86 [4.06, 7.31] | 8.10 [6.37, 12.18] |
| *Ozomelis* | 4.19 [2.95, 5.18] | 3.71 [2.63, 4.86] | – |
| *Ozomelis* group | 11.42 [10.94, 11.98] | 10.85 [10.57, 11.19] | – |
| *Pectiantia* group | 11.34 [10.84, 12.14] | 10.34 [10.07, 10.61] | – |
| *Tiarella* | 5.15 [3.89, 6.08] | 5.51 [3.83, 7.34] | – |
| *Tolmiea* | 4.48 [1.49, 6.40] | 5.5 [3.14, 8.08] | 3.6155 [0.7, 5.57] |

**Table S3.** Summary of inferred chloroplast capture event clades and dates, reflective of Fig. 5. In the case of eastern species of *Heuchera* (subsects. *Heuchera* and *Villosae*), there is complex cytonuclear discordance at the population level. In the absence of sufficient population-level sampling these have been marked as "unresolved."

| **Putative donor** | **Putative recipient** | **Date (chloroplast analysis MRCA)** | **First report** |
|---|---|---|---|
| MRCA(*Mitella*) | MRCA(*Heuchera caespitosa*, *H. elegans*) | 3.17 | Folk et al. 2017 |
| MRCA(*Lithophragma heterophyllum*, *L. bolanderi*) | MRCA(*Tolmiea*) | 3.62 | This study |
| *Tiarella* | *Holochloa* group | 4.00 | Soltis et al. 1991 (but resolved differently here, with *T. cordifolia* treated below) |
| *Heuchera micrantha* | *H. wootonii* | 2.86 | Folk et al. 2017 |
| *H. cylindrica* (population treated historically as "*H. saxicola*") | *H. grossulariifolia* 4x | 1.07 | Folk et al. 2017 (sampled but unresolved in Soltis et al. 1991) |
| *H. cylindrica* ("*H. saxicola*"; different population from above) | *H. grossulariifolia* 2x | 0.91 | Folk et al. 2017 (sampled but unresolved in Soltis et al. 1991) |
| *H. cylindrica* | *H. parvifolia* (populations historically treated as "*H. duranii*") | 1.66 | Folk et al. 2017 |
| *H. cylindrica* | *Ozomelis* | 2.59 | Folk et al. 2017 (sampled but unresolved in Soltis et al. 1991) |
| *Ozomelis* | *H. micrantha* (northern populations treated as vars. *diversifolia, macropetala*) | 1.39 | Folk et al. 2017 (sampled but unresolved in Soltis et al. 1991) |
| *H. micrantha* var. *macropetala* | *H. parvifolia* var. *utahensis* | 0.82 | Folk et al. 2017 |
| *H. micrantha* | *H. glabra* | 0.31 | Folk et al. 2017 (sampled but unresolved in Soltis et al. 1991) |
| Unresolved member of *Heuchera* subsect. *Villosae* | *Tiarella cordifolia* | 1.38 | Folk et al. 2017 (Sanger analysis) |
| Unresolved member of *Heuchera* subsect. *Villosae* | MRCA(*Heuchera* subsect. *Heuchera*) | 1.72 | Folk et al. 2017 |
| Unresolved member of *Heuchera* subsect. *Villosae* | Unresolved members of *Heuchera* subsect. *Heuchera* | 1.52 | Folk et al. 2017 |
| Unresolved member of *Heuchera* subsect. *Villosae* | Unresolved members of *Heuchera* subsect. *Heuchera* | 0.82 | Folk et al. 2017 |
| MRCA(*Asimitellaria japonica*, *A. yoshinagae*) | *Asimitellaria yamatoensis* | 2.31 | Okuyama et al. 2005 |
| *A. stylosa* | *A. acerina* | 1.15 | Okuyama et al. 2005 |

**Table S4.** Comparison of monophyly support among recent multilocus phylogenetic studies, focusing on named clades, with two additional major subclades for each of *Asimitellaria* and *Lithophragma*. "—" means the clade was not recovered; "X" means the sampling was not sufficient to test the presence of the clade.

| **Clade** | **This study, ASTRAL (LPP)** | **This study, concatenation (bootstrap)** | **Folk et al. 2018c** | **Folk et al. 2017, ASTRAL (multilocus bootstrap)** | **Folk et al. 2017, gene-partitioned concatenation (bootstrap)** | **Folk and Freudenstein 2014 total evidence, concatenated (ML bootstrap)** | **Okuyama et al. 2012 concatenated (bootstrap; subgenome A, subgenome B for *Asimitellaria*)** |
|---|---|---|---|---|---|---|---|
| *Asimitellaria* | 1 | 100 | 100 | X | X | X | 100 |
| MRCA(*Asimitellaria formosana*, *A. japonica*) | 1 | 100 | X | X | X | X | 99, 100 |
| MRCA(*Asimitellaria stylosa*, *A. furusei*) | 1 | 100 | 100 | X | X | X | 82, 100 |
| *Brewerimitella* | 1 | 100 | 100 | X | X | X | 100 |
| *Heuchera* | 1 | 100 | 100 | 100 | 100 | 100 | 100, 100 |
| *Heuchera* sect. *Heuchera* | 1 | 100 | 100 | 100 | 100 | 100 | X |
| *Heuchera* sect. *Holochloa* (incl. *H. glabra*) | 0.94 | — | 85 | 100 | 95 | — | X |
| *Heuchera* sect. *Rhodoheuchera* | 0.69 | — | < 50 | 100 | 27 | 86 | X |
| *Heuchera* subsect. *Cylindricae* | 1 | 100 | 100 | 100 | 100 | — | X |
| *Heuchera* subsect. *Elegantes* | 0.9 | — | 71 | 98 | — | — | X |
| *Heuchera* subsect. *Heuchera* | 1 | 96 | 100 | 100 | 100 | 73 | X |
| *Heuchera* subsect. *Micranthae* | 1 | 100 | 100 | 100 | 100 | 100 | X |

| | | | | | | | |
|---|---|---|---|---|---|---|---|
| *Heuchera* subsect. *Novomexicanae* | 1 | 100 | 100 | 100 | 100 | — | X |
| *Heuchera* subsect. *Villosae* | 1 | 100 | 100 | 100 | 100 | 100 | X |
| *Lithophragma* | 1 | 100 | X | X | X | X | X |
| MRCA(*Lithophragma heterophyllum, L. maximum*) | 1 | 92 | X | X | X | X | X |
| MRCA(*Lithophragma parviflorum, L. tenellum*) | 1 | 92 | X | X | X | X | 99 |
| *Mitella* | | | | | | | 97 |
| *Ozomelis* | 1 | 100 | X | X | X | X | X |
| *Ozomelis* group | 1 | 100 | 100 | X | X | 52 | 90 |
| *Pectiantia* group | 0.65 | 100 | 95 | X | X | — | — |
| *Tiarella* | 1 | 100 | X | X | X | X | 100 |
| *Tolmiea* | 1 | 1 | 100 | X | X | X | X |
| Tribe Heuchereae | 1 | 100 | 100 | X | X | 100 | 100 |

**Table S5.** Comparison of ancestral niche reconstruction results between the new method and two previously developed methods. Results represent BIO1 (mean annual temperature), formatted as center/range. Centers report niche centroid as measured by each of these methods: max bin height (histogram method), maximum likelihood estimate (Phyloclim), and posterior median (Ambitus). Ranges report niche breadth as measured by each of these methods: range from min to max bin with at least 1% probability (histogram method) and 95% credibility interval max - min (Ambitus); note Phyloclim does not directly implement ancestral node estimates of niche breadth. Nodes reported represent the tribe and all non-monotypic genera.

| **Node** | **Histogram method** | **Phyloclim (Heibl 2011)** | **Ambitus (Folk et al. 2017)** |
|---|---|---|---|
| MRCA (tribe Heuchereae) | 9.844/18.564 °C | 9.206 °C | 7.453/9.252 °C |
| MRCA(genus *Heuchera*) | 10.228/18.564 °C | 8.942 °C | 7.833/8.858 °C |
| MRCA(genus *Lithophragma*) | 12.028/17.472 °C | 11.56 °C | 10.256/7.982 °C |
| MRCA(genus *Asimitellaria*) | 12.028/13.114 °C | 10.389 °C | 11.238/6.867 °C |
| MRCA(genus *Mitella*) | 7.66/21.84 °C | 6.683 °C | 5.793/11.227 °C |
| MRCA(genus *Brewerimitella*) | 9.844/16.38 °C | 5.999 °C | 6.54/10.25 °C |
| MRCA(genus *Tiarella*) | 5.476/18.564 °C | 8.162 °C | 7.563/11.339 °C |
| MRCA(genus *Ozomelis*) | 6.568/15.288 °C | 5.38 °C | 5.076/10.252 °C |
